# Vesicular acidification modulates the synaptic current: a hybrid diffusion–reaction model analysis

**DOI:** 10.64898/2026.04.23.720425

**Authors:** R. Bar-on, I.H. Greger, D. Holcman

## Abstract

Glutamate synaptic vesicles co-release protons, producing a brief acidification of the synaptic cleft that could modulate AMPA receptor (AMPARs) operation. To evaluate the extent of receptor acidification, we develop a diffusion–reaction model that couples vesicle-evoked proton and glutamate transients to AMPAR dynamics. Our simulations reveal that the rapid diffusion of protons and glutamate within the flat-cylindrical synaptic cleft leads to a mixture of protonated, singly glutamate-bound and doubly glutamate-bound AMPARs. We studied four postsynaptic AMPAR distributions - uniform disk, sub-disk, Gaussian cluster, and point-like cluster - and showed a ∼ 50% increase in the number of acidified receptors when AMPARs are clustered on the postsynaptic cleft compared to a uniform arrangement. We further explored the impact of pH revealing that at acidic conditions (pH ∼ 5), approximately 80–90% of open receptors are non-acidified, whereas under strongly acidic conditions (pH ∼ 3), about 80–90% of open receptors exist in the protonated form. Finally, we explored how acidification modulates AMPARs during paired-pulse stimulation, a measure of short-term synaptic depression. While the presence of protons does not markedly alter the overall trends, acidified receptor states reduce the occupancies of their neutral counterparts by roughly 10–15%, indicating a mild redistribution toward protonated receptor conformations. To conclude, our model suggests that AMPAR protonation can influence the synaptic current, and we predict that this effect is determined primarily by the dissociation kinetics of glutamate and protons from AMPAR.

## 1 Introduction

Fast excitatory glutamatergic synaptic transmission relies on highly specialized, submicron contacts between neurons that enable neurotransmiter release from vesicles [1, 2, 3]. Inside the sub-micron cleft geometry, glutamate diffuse to activate postsynaptic *α*-amino-3-hydroxy-5-methyl-4-isoxazolepropionic acid receptors (AMPARs) [4, 5, 6, 7]. The number of activated AMPARs is a key determinant of the postsynaptic current. An increase in this number underlies long-term potentiation (LTP) of synaptic transmission, a process central to learning and memory [8, 9]. The organization of the synaptic cleft geometry could be determinant of the postsynaptic current [10, 11, 12, 13, 14, 15, 16, 17] through trans-synaptic organization of location of vesicular release and AMPAR clustering [18, 19]. Thus the magnitude and temporal profile of the postsynaptic response are not determined solely by the biophysical receptor properties, but emerge from the interplay of presynaptic, synaptic, and postsynaptic factors, such as the location of vesicular release, the geometry of the cleft, the spatial distribution, and kinetic properties of each AMPAR type and finally the presence of surrounding glial processes [12, 20, 21, 22, 23, 24, 25, 26].

Computational and theoretical studies that incorporate synaptic geometries and diffusion have shown that variations in these parameters can affect the time course and amplitude of AMPAR-mediated responses, together with receptor binding and gating kinetics [12, 10, 27, 13, 23, 28, 29]indicating that the differences in synaptic morphology and AMPAR distribution could contribute to the heterogeneity of excitatory postsynaptic current (EPSC) kinetics across central synapses [12, 23, 30]. Such a process depends not only on the biophysical properties of AMPAR subunits [4], but also on receptor trafficking [31, 32], that can enter and exit the post-synaptic density (PSD), thus influencing the post-synaptic current responses across a time of few tens of ms. Trafficking modifies the synaptic strength across time and in addition, trial-to-trial variability depends also on the alignment (multiple nano-columns) of the active zone with AMPAR clusters [33, 10, 34] and the postsynaptic density (PSD)/perisynaptic organization. While trafficking can partially rescue short-term depression [35, 19], structural remodeling of AMPAR can modulate the post-synpatic current [36]. Moreover, the PSD morphology and release location can influence the peak post-synaptic current and variability, as evaluated in multi-conductance AMPAR models [11], leading to an optimal PSD size that minimizes response variance—a geometry-based form of plasticity. In the present manuscript, we study how the pH dynamics can influence AMPARs and thus the post-synaptic current. It was recently reported that synaptic vesicular release produces brief acidification transients with pH ≈ 5.3–5.7 in the cleft before alkalinization, suggesting that protons can modulate receptors [37, 38], which is essential for neurotransmitter loading and quantal release. Vesicular acidification is maintained by vacuolar-type H^+^-ATPases (V-ATPases), which actively pump protons into the vesicle lumen, generating both a proton electrochemical gradient and a low intravesicular pH that drives glutamate uptake via vesicular glutamate transporters [39]. Measurements using pH-sensitive fluorescent reporters have shown that the resting pH inside synaptic vesicles is tightly regulated [40]. Upon exocytosis, this acidic content is transiently released into the synaptic cleft, producing brief proton transients that can locally modulate synaptic signaling [41]. These observations establish vesicular release as a physiologically relevant source of synaptic acidification and constrain the expected proton load associated with a single vesicular event [42, 43, 44]. A GluA2/TARP-*γ*2 cryo-EM structure determined at acidic pH revealed that acidification can tune AMPAR structure, gating kinetics and diffusion at the synapse [37, 38]: protonation targets a GluA2-specific N-terminal domain (NTD) interdimer interface centered on histidine H208, rupturing the compact NTD tier and propagating rearrangements across the extra-cellular region [37, 38], as also observed with the GluA2 F231A mutation [45, 36]. Consequently, low pH reduces peak and steady-state currents by lowering the open probability without changing unitary conductance [46, 47, 42], speeds desensitization entry, slows recovery, accelerates deactivation, and right-shifts the glutamate dose–response (EC50 ∼ 0.27 → 1.84 mM for GluA2+TARP *γ*2) [37, 38]. Another consequence of acidification is to increase synaptic receptor mobility [37, 38], linking gating to trafficking. Our goal here is to integrate the proton allostery change into a coarse-grained synaptic model to evaluate the consequence of acidification on the synaptic current. The modulation of AMPAR gating by extracellular protons has been investigated in earlier electrophysiological and structural studies. Acidic pH was shown to reduce AMPAR peak currents primarily by enhancing receptor desensitization and accelerating entry into desensitized states, without altering unitary conductance [46]. Consistently, proton-induced inhibition was found to shift the glutamate dose–response and to interact with receptor allosteric modulators, which can partially rescue proton-driven desensitization [47]. More recently, structural and functional work has demonstrated that recovery from desensitization in GluA2-containing AMPARs critically depends on the integrity of the N-terminal domain (NTD) interdimer interface [38], a region now known to be sensitive to protonation-induced rearrangements. Together, these studies establish extracellular protons as key modulators of AMPAR gating kinetics and provide a mechanistic framework for interpreting proton-dependent effects on synaptic responses.

The manuscript is organized as follows: we first develop a reaction–diffusion equation in a cylindrical geometry to quantify the transient proton dynamics and estimate the binding to AMPARs. In the result section, we first quantify proton diffusion and clearance within the synaptic cleft and determine how the spatial organization of AMPARs and vesicular release location shapes receptor acidification following a single vesicular event. We then analyze the coupled proton–glutamate binding dynamics of AMPARs to characterize the temporal evolution of receptor states and identify the regimes in which protonation alters glutamate-driven activation. Next, we examine how variations in proton load and kinetic rate constants influence receptor occupancy and the competition between protonated and non-protonated pathways. Finally, we extend our analysis to paired-pulse stimulation, assessing how transient acidification modulates recovery and desensitization across pH ranges and kinetic regimes. Together, our model and simulations allows to evaluate how pH transients reshape AMPAR signaling during synaptic activity and this effect depend on backward rate constant that should be experimentally measured.

## 2 Results

### 2.1 Quantifying AMPAR acidification following a single vesicular release

To quantify the number of AMPAR protonated following a single vesicular event [37, 38], we develop here a simulation involving the coupling between reaction and diffusion (Method Section) based on proton diffusion in the synaptic cleft, that could interact with AMPARs located on the post-synaptic side. Protons with concentration *u*(*r, t*) eq. (2) are released uniformly (eq. (3)) from a sub-disk of radius 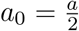 at the top surface of the cleft (Fig. 1A-C) with height *L* = 20 nm (Table 2.1).

**Table 1:** Parameters used in the simulations. In the upper part of the table, parameters have been kept fixed during all simulations. Middle: parameters have been chosen at order of magnitude with respect to *k*_des_ and *k*_rec_. Lower part of the table: parameters were chosen as reasonable values based on time scales reported in [48, 49, 50, 51].

| Parameter | Value | Description |
| --- | --- | --- |
| $a$ | $0.5 \mu\text{m}$ [52] | Synaptic disk radius |
| $a_0$ | $a/2 = 0.25 \mu\text{m}$ | Sub-disk radius |
| $L$ | $0.02 \mu\text{m}$ [10] | Cleft height |
| $N_{\text{AMPAR}}$ | 50 [52] | Total number of receptors |
| $\sigma_{\text{Dirac}}$ | $0.01 a$ | Width for sharp (Dirac-like) Gaussian |
| $\sigma_{\text{Gauss}}$ | $0.10 a$ | Width for broader Gaussian |
| $D$ | $500 \mu\text{m}^2/\text{s}$ | Proton diffusion coefficient |
| $D_g$ | $500 \mu\text{m}^2/\text{s}$ [53] | Glutamate diffusion coefficient |
| $b$ | $0.001 \mu\text{m}$ [9] | Capture AMPAR radius for binding |
| $k_{-1}$ | $10^4 \text{s}^{-1}$ | Proton unbinding rate |
| $k_{-2}$ | $10^5 \text{s}^{-1}$ | First glutamate unbinding rate (protonated state) |
| $k_{-3}$ | $5 \times 10^6 \text{s}^{-1}$ | Second glutamate unbinding rate (protonated state) |
| $k_{-4}$ | $10^5 \text{s}^{-1}$ | First glutamate unbinding rate (non-protonated state) |
| $k_{-5}$ | $5 \times 10^6 \text{s}^{-1}$ | Second glutamate unbinding rate (non-protonated state) |
| $k_{-1}$ | $200 \text{s}^{-1}$ | Proton unbinding rate |
| $k_{-2}$ | $200 \text{s}^{-1}$ | First glutamate unbinding rate (protonated state) |
| $k_{-3}$ | $200 \text{s}^{-1}$ | Second glutamate unbinding rate (protonated state) |
| $k_{-4}$ | $200 \text{s}^{-1}$ | First glutamate unbinding rate (non-protonated state) |
| $k_{-5}$ | $200 \text{s}^{-1}$ | Second glutamate unbinding rate (non-protonated state) |
| $\langle k_{\text{des}}^H(r, t) \rangle$ | $71.6 \text{s}^{-1}$ [37] | Proton-modulated desensitization rate |
| $\langle k_{\text{rec}}^H(r, t) \rangle$ | $63.8 \text{s}^{-1}$ [37] | Proton-modulated recovery rate |

**Figure 1:**
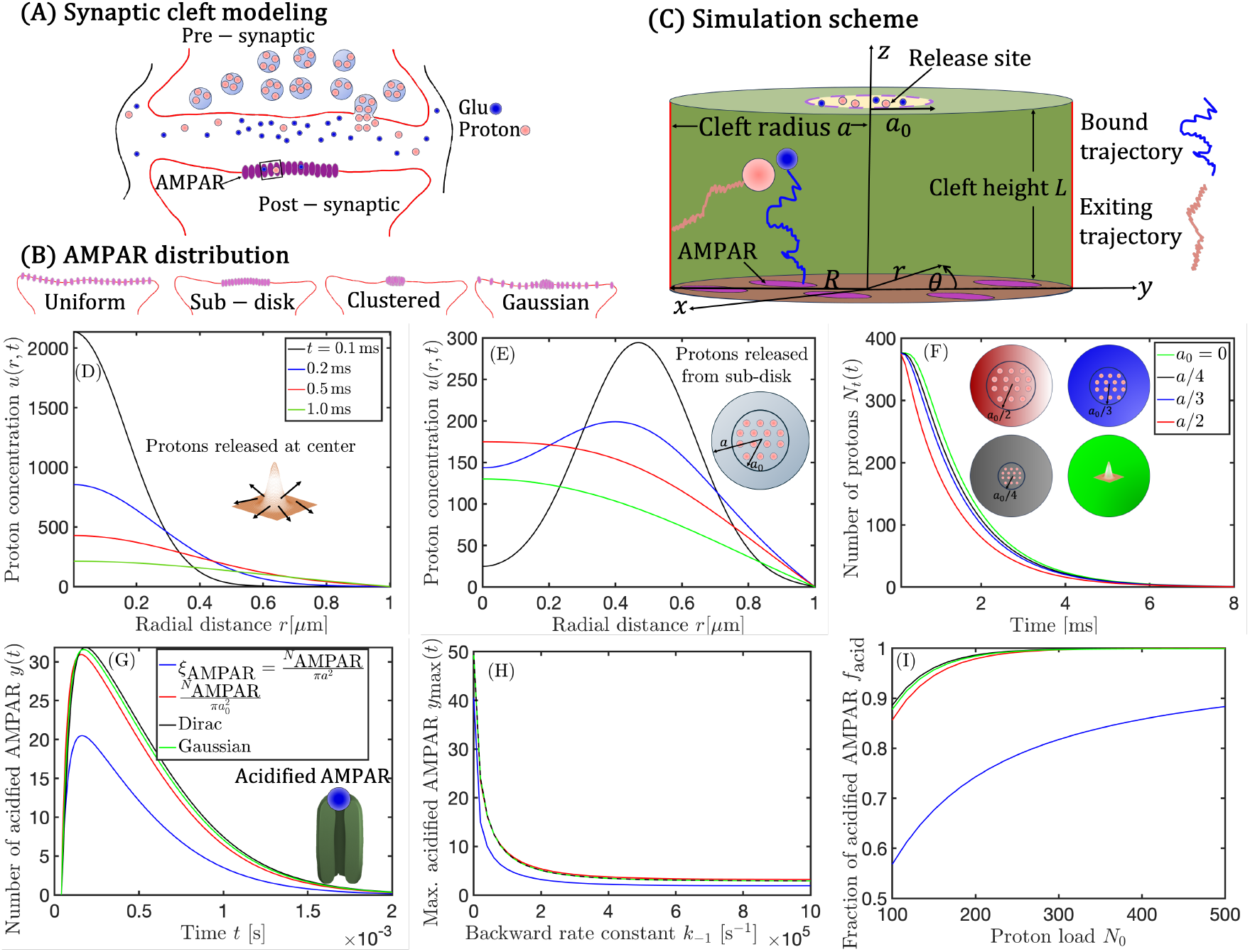
AMPAR acidification in a synaptic cleft following a single vesicular event. **(A)** Schematic representation of proton and glutamate dynamics within the synaptic cleft following vesicular release. Glutamate (pink) and protons (blue) diffuse throughout the cleft and interact with AMPARs (magenta) distributed uniformly on a sub-disk of the postsynaptic membrane. **(B)** Alternative AMPAR spatial distributions on the postsynaptic terminal: uniform across the full cleft, uniform on a sub-disk, clustered, and Gaussian-like. **(C)** Computational domain Ω of the synaptic cleft used in the numerical simulations. Proton (blue) and glutamate (pink) trajectories originate from a sub-disk of radius *a*_0_ centered at *r* = 0. AMPARs (magenta) are distributed within a disk of radius *R* on the postsynaptic surface. **(D)** Proton radial concentration *u*(*r, t*) at various time points (*t* = 0.1, 0.2, 0.5, 1 ms) following an initial release at the center of the presynaptic terminal (*N*_0_ = 376). Simulation parameters are listed in Table 2.1. **(E)** Same as in (D), except the initial release occurs from a narrow annulus of radius *a*_0_ = *a/*2. **(F)** Temporal evolution of the total number of protons *N* (*t*) for four release configurations: (a) centered, (b) *a*_0_ = *a/*2, (c) *a*_0_ = *a/*3, and (d) *a*_0_ = *a/*4. **(G)** Time course of acidified AMPARs *y*(*t*) = [H^+^A] for a backward rate constant *k*_−1_ = 10^4^ s^−1^, comparing the four receptor distributions (see Fig. 1(B)). **(H)** Maximum number of acidified AMPARs *y*_max_ as a function of the backward rate *k*_−1_ for the four distributions defined in (B). **(I)** Fraction *f*_acid_ of acidified AMPARs (i.e., receptors acidified at least once during the simulation) as a function of the initial proton load *N*_0_. The corresponding cleft pH is obtained from 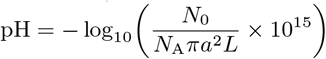, where *N*_A_ is Avogadro’s constant, *a* is the cleft radius, and *L* its height.

To evaluate the rate of proton escape, we look at snap shots at short-time steps *t* = 0.1, 0.2, 0.5, and 1.0 ms, the simulations show that most protons are removed after 1ms, whether the release occurs on the center or in a sub-disk of a radius 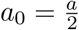 (Fig. 1D-E). The total number *N*_total_(*t*) of protons in the cleft, decays after a few ms, independently of the initial release radii: *a*_0_ = 0 (green), *a*_0_ = *a/*4 (black), *a*_0_ = *a/*3 (blue), and *a*_0_ = *a/*2 (red) (Fig. 1F). To quantify the acidified AMPAR concentration *y*(*t*) vs time, for the four spatial distributions of AMPAR, we simulated proton release from a sub-disk (Fig. 1(G)). When receptors are uniformly distributed over the entire disk, the maximum number of AMPAR is 20 (from a total of 50), whereas the other distributions result in peaks 30. Since the reverse reaction rate constant *k*_−1_ of an acidified AMPAR is unknown, we vary it across several orders of magnitude, ranging from 10^2^ s^−1^ to 10^6^ s^−1^ (Fig. 1(H)). *y*_max_(*t*) varies between 3 to approximately 50 (all receptors considered to be acidified), as the backward rate constant *k*_−1_ decreases.

To test the effect of proton load/pH on the number of acidified AMPAR, we calculated the fraction *f*_acid_ of acidified AMPAR vs the initial proton number *N*_0_ (Fig. 1(I)): for low *N*_0_ values (*N*_0_ *<* 300), *f*_acid_ is highly sensitive to the initial proton concentration, with noticeable differences between some of the AMPAR spatial distributions. The uniform distribution over the entire disk (blue curve) shows the lowest *f*_acid_. The sub-disk, centered release and Gaussian distributions (red and black curves, respectively) yield higher *f*_acid_, approaching 1 more rapidly, due to the localized nature of proton release.

For high *N*_0_ values (*N*_0_ *>* 300), the sub-disk, centered and Gaussian distribution lead to *f*_acid_ ∼ 1, whereas the uniform distribution conformation produces similar effect for higher *N*_0_. To conclude, this result indicates a saturation effect, where further increase in *N*_0_ results in minimal additional acidification, as the AMPAR sites are already maximally occupied.

### 2.2 Dynamics of proton and glutamate bound to AMPARs

Having characterized proton diffusion and receptor acidification, we next examined the coupled proton–glutamate dynamics governing AMPAR activation. Our goal is to quantify how each receptor state evolves in time following a single vesicular release, and how spatial receptor organization influences these transitions. Figure 2 summarizes the temporal evolution of the six receptor populations defined in the reaction scheme, simulated for four spatial distributions of AMPARs: uniform over the full postsynaptic disk, uniform over a sub-disk, Gaussian-like, and centrally localized.

**Figure 2:**
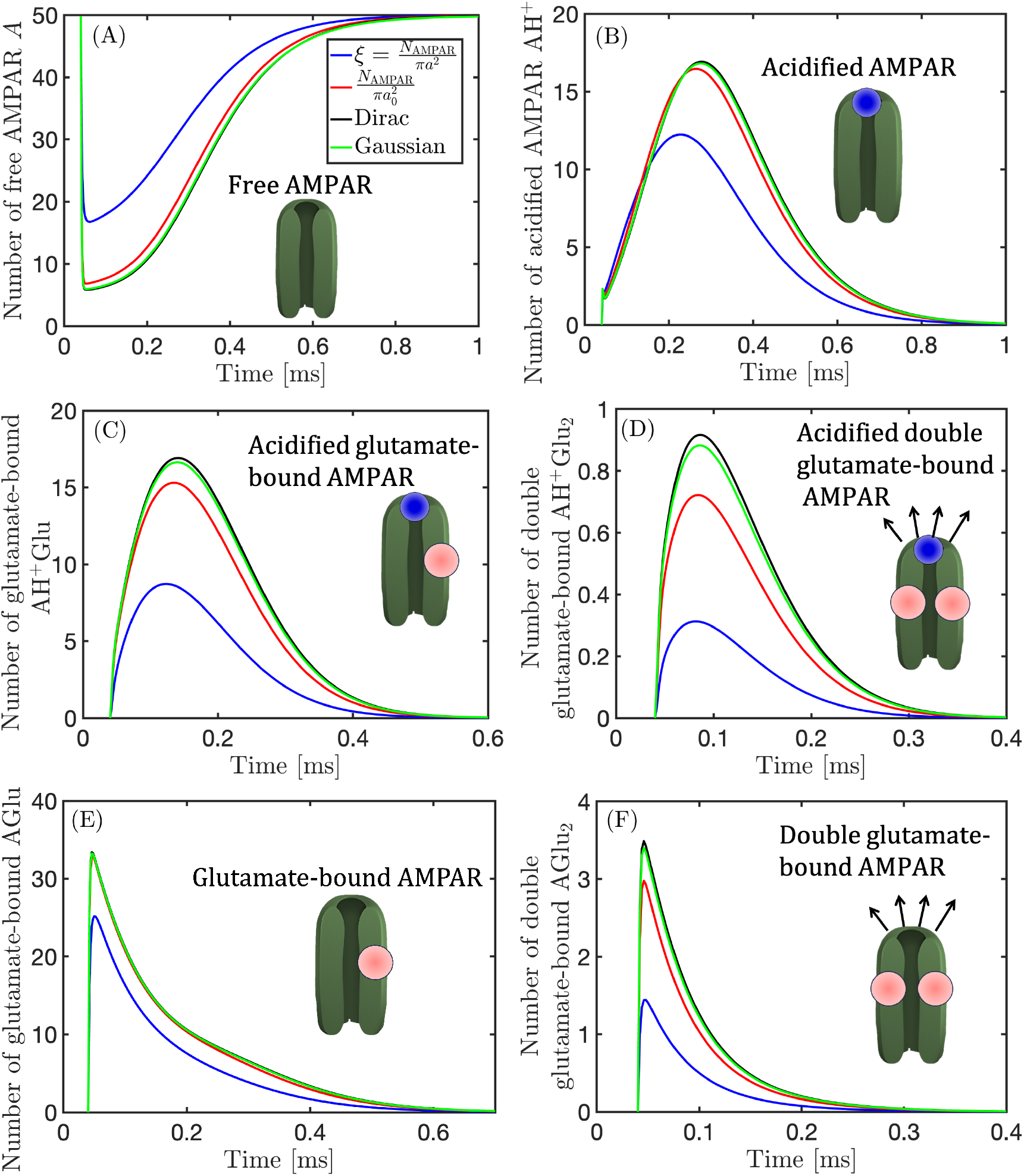
Dynamics of bound AMPAR species for four spatial receptor distributions. **(A)** Time course of free receptors [A]. Simulation parameters are listed in Table 2.1. **(B)** Acidified receptors [AH^+^]. **(C)** Acidified receptors bound to a single glutamate [AH^+^Glu]. **(D)** Acidified receptors bound to two glutamates [AH^+^Glu_2_]. **(E)** Non-acidified receptors bound to one glutamate [AGlu]. **(F)** Non-acidified receptors bound to two glutamates [AGlu_2_]. Each panel compares the four receptor distributions defined in Fig. 1B: uniform full-disk, uniform sub-disk, Clustered (modeled as Dirac-like Gaussian), and Gaussian

Immediately after release, the fraction of free receptors [A] (Fig. 2A) drops sharply as binding events occur, followed by a gradual recovery once ligands dissociate or diffuse away. The rapid initial depletion (sub-millisecond) indicates that both proton and glutamate binding are fast relative to unbinding. Broader receptor distributions sustain slightly higher free-receptor levels at early times, reflecting reduced local crowding near the release center.

The solely acidified state [AH^+^] (Fig. 2B) rises transiently to a maximum of ∼ 12– 17 receptors within 2 *×* 10^−4^ s, then decays as protons unbind or are displaced by glutamate. This brief peak underscores the reversibility of protonation and its competition with subsequent glutamate binding, which dominates at later stages.

Following protonation, some receptors acquire glutamate sequentially. The intermediate acidified–glutamate state [AH^+^Glu] (Fig. 2C) emerges rapidly, reaching ∼ 10–18 receptors at ∼ 1.5 *×* 10^−4^ s. Only a small fraction proceeds to the fully bound acidified–diglutamate state [AH^+^Glu_2_] (Fig. 2D), which peaks at ∼ 1 receptor, consistent with the short glutamate lifetime in the cleft and competition with non-protonated receptors.

The singly bound [AGlu] (Fig. 2E) and doubly bound [AGlu_2_] (Fig. 2F) populations rise almost instantaneously, with maxima around 25–33 and 2–5 receptors, respectively. Since backward reaction rate constants *k*_−1_, *k*_−2_, *k*_−3_ of a proton, and *k*_−4_, *k*_−5_ of a glutamate bound to AMPAR are unknowns, we decided to explore various range of parameters under which the reaction rates are stable (Fig. S1). The ratio 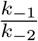 is almost constant for 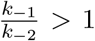. However for 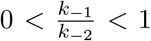, the max number of AH^+^ decreases from 40 to 5 (Fig. S1A). Similarly, The ratio 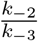 is almost constant for 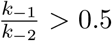. However for 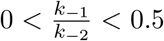, the max number of AH^+^Glu decreases from 40 to 5 (Fig. S1B). Similarly, we explored the ratio 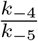 and obtained comparable results (Fig. S1C).

Finally, we explore the range for *k*_−3_ (resp. *k*_−5_) that represents the dissociation of glutamate from AH^+^Glu_2_ (resp. AGlu_2_), motivated by the single chemical pathway. In both cases, for a backward constant larger than 10^5^, we observe no much changes Fig. S1D-E). We note that the effect of spatial distribution of the AMPAR is marginal as shown by the different color curves.

To assess how receptor occupancy evolves over time, we next analyzed the percentage of protonated and AMPAR glutamate-bound in the limiting case with no backward reaction by setting *k*_−1_ = *k*_−2_ = *k*_−3_ = *k*_−4_ = *k*_−5_ = 0 (Fig. S2), across the four spatial receptor distributions *ξ*_AMPAR_(*r*) (Fig. S2). Our results suggest that the effect of acidification on AMPAR states is minor as only 3% − 4% of all receptors go to AH^+^ state.

### 2.3 Impact of acidification on bound AMPAR

We next explore how the initial proton load, *N*_0_, influences the occupancy of each receptor state, for the uniform distribution of AMPAR on the post synaptic cleft. By scanning values from *N*_0_ = 100 up to *N*_0_ = 10^4^, we assess how proton availability modulates both the amplitude and temporal profile of the binding dynamics (Fig. 3A–F). Physiologically, the proton content released by a synaptic vesicle is constrained by the intravesicular pH, which is tightly regulated and typically lies around pH ≈ 5.3–5.7. Nevertheless, in the present work we deliberately explore a broader range of initial proton loads, including values corresponding to more acidic conditions. This choice is motivated by the mathematical nature of the model, which is designed to probe the sensitivity of AMPAR binding dynamics to proton availability and kinetic competition across regimes that may extend beyond strict physiological conditions. In particular, considering lower effective pH values allows us to delineate limiting behaviors of the coupled proton–glutamate system and to identify thresholds at which protonation qualitatively reshapes receptor occupancy. We note that at sufficiently low pH, such as pH ∼ 3, structural studies indicate that the AMPAR N-terminal domain undergoes pronounced rearrangements and loss of compact organization [37, 38]. While such conditions are unlikely to occur in vivo within the synaptic cleft, their inclusion here serves as a controlled theoretical exploration that helps clarify the role of proton-driven allosteric modulation in shaping synaptic responses. For the free receptor pool [A], increasing *N*_0_ leads to progressively deeper initial minima (Fig. 3A). With only 100 protons the minimum remains near 18 receptors, while for *N*_0_ = 10^4^ almost all receptors are depleted to values as low as ∼ 2 − 3 within the first 0.1 ms, before rebounding as bound states decay. The acidified-only state [AH^+^] exhibits sharp, dose-dependent transients (Fig. 3B). For *N*_0_ = 100 the peak barely exceeds ∼ 5 receptors, whereas for *N*_0_ = 10^4^ the maximum surpasses 30. In all cases the response is short-lived (*<* 1 ms). The singly glutamate-bound acidified complex [AH^+^Glu] grows monotonically with *N*_0_, with maxima ranging from ∼ 3 − 4 receptors at low proton counts to ∼ 26 − 27 at high counts (Fig. 3C). The doubly glutamate-bound acidified complex [AH^+^Glu_2_] remains rarer (Fig. 3D), but still increases from ∼ 0 to ∼ 2 receptors as *N*_0_ is raised. Glutamate-dominated pathways behave in the opposite manner. The non-protonated singly bound state [AGlu] (Fig. 3E) reaches its maximum almost immediately after release, but the amplitude decreases systematically with increasing proton number. For a low proton load (*N*_0_ = 100), peaks approach ∼ 28 receptors, while for *N*_0_ = 10^4^ the maximum is reduced to only ∼ 7. The doubly glutamate-bound state [AGlu_2_], follows the same trend (Fig. 3F): from ∼ 2 receptors at low *N*_0_, the peak shrinks to below 0.5 as proton availability increases. This inverse relationship reflects competition between protons and glutamate: higher proton concentrations transiently trap receptors in acidified or mixed states, leaving fewer available to proceed along the purely glutamate-driven pathway.

**Figure 3:**
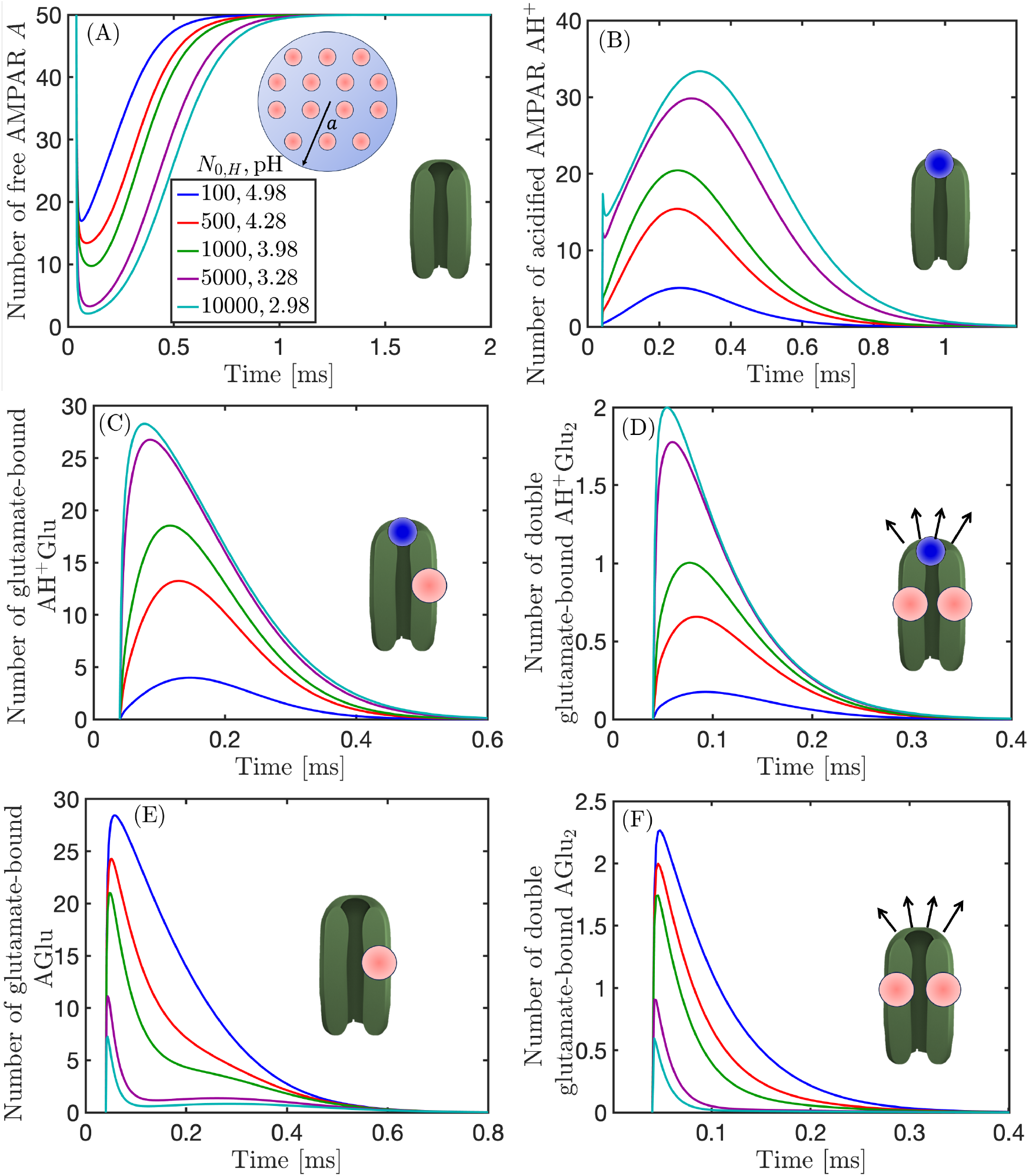
Dynamics of AMPAR species for increasing proton loads *N*_0_, for a uniform release. **(A)** Free receptors [A]. **(B)** Acidified receptors [AH^+^]. **(C)** Acidified receptors bound to a single glutamate [AH^+^Glu]. **(D)** Acidified receptors bound to two glutamates [AH^+^Glu_2_]. **(E)** Non-acidified receptors bound to a single glutamate [AGlu]. **(F)** Non-acidified receptors bound to two glutamates [AGlu_2_]. Curves correspond to increasing proton release numbers: *N*_0_ = 100 (blue), 500 (red), 1000 (dark green), 5000 (purple), and 10000 (teal).

#### 2.3.1 Proton initial number impact on the maximum of AMPAR-bound species

We next quantify how the maximum number of receptors attained in each state depends on the proton load *N*_0_, as summarized in Fig. 4. For the acidified-only state [AH^+^], the peak occupancy grows monotonically with *N*_0_, rising from ∼ 5 − 10 (ranging from uniform distribution to sub-disk and centered distributions) receptors at *N*_0_ = 100 to nearly ∼ 30 − 35 at *N*_0_ = 10^4^ (Fig. 4A). A similar trend holds for the mixed state [AH^+^Glu], whose maxima increase from ∼ 5 − 10 receptors at low *N*_0_ to ∼ 20–30 at the highest values (Fig. 4B). The doubly glutamate-bound acidified complex [AH^+^Glu_2_] shows the same monotonic increase but on a smaller scale (Fig. 4C) of ∼ 1 − 3 receptors.

**Figure 4:**
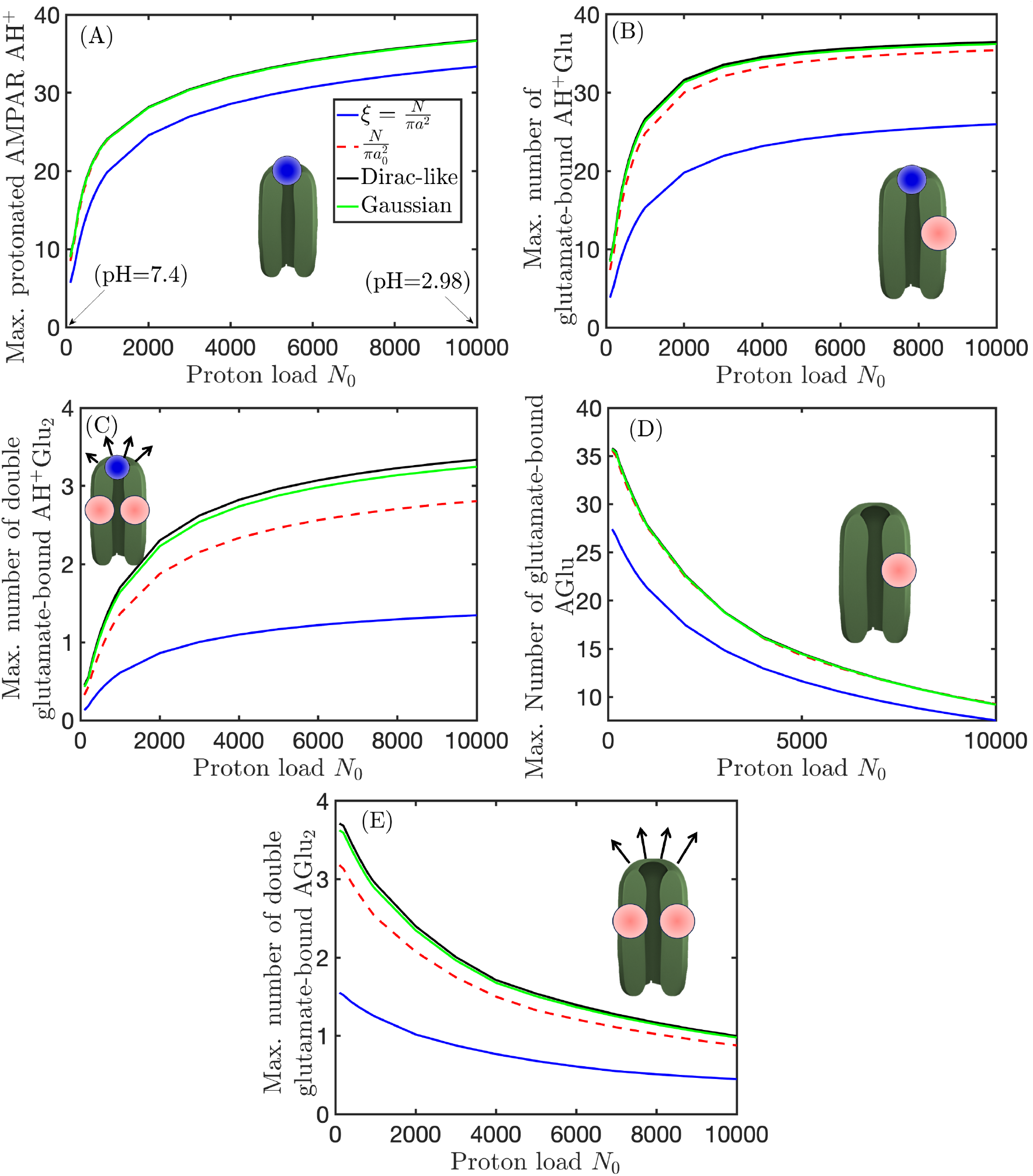
Maximum number of AMPAR states as a function of the initial proton load *N*_0_, for the four spatial receptor distributions. **(A)** Acidified receptors [AH^+^]. **(B)** Acidified receptors bound to a single glutamate molecule [AH^+^Glu]. **(C)** Acidified receptors bound to two glutamate molecules [AH^+^Glu_2_]. **(D)** Non-acidified receptors bound to one glutamate [AGlu]. **(E)** Non-acidified receptors bound to two glutamates [AGlu_2_].

In contrast, glutamate-dominated pathways are suppressed by increasing proton load. The non-protonated singly bound state [AGlu] decreases with *N*_0_, dropping from ∼ 27 − 36 receptors at *N*_0_ = 100 to ∼ 0 − 10 at *N*_0_ = 10^4^ (Fig. 4D). The doubly glutamate-bound state [AGlu_2_], representing the open-channel precursor, mirrors this suppression (Fig. 4E), declining from ∼ 2 − 4 receptors for the clustered distribution to around unity as *N*_0_ increases.

### 2.4 Paired–pulse dynamics of AMPAR states under glutamate release only (Δ = 50 ms)

We next analyzed how a second, identical glutamate release delivered after a delay of Δ = 50 ms reshapes receptor occupancy and recovery, under conditions where only glutamate-driven reactions are active (no protonation). To assess whether there could be an even slight difference between the two peaks, we use comparable backward rate constants to AMPAR desensitization and recovery rates reported [37, 38]), See Table 2.1. Paired–pulse responses are not determined solely by intrinsic AMPAR binding, desensitization, and recovery kinetics, but can also be influenced by the lateral mobility of receptors at the postsynaptic membrane. In particular, AMPAR surface diffusion enables rapid exchange between synaptic and extrasynaptic receptor pools on the tens-of-milliseconds timescale, thereby contributing to recovery from synaptic depression following repeated stimulation [35]. In the present analysis, receptor mobility is not explicitly modeled, allowing us to isolate the contribution of kinetic processes to paired–pulse dynamics under glutamate-only and glutamate-plus-proton conditions.

Following each release, the free receptor population [A] (Fig. 5(A)) exhibits nearly identical depletion–recovery transients for all receptor spatial distributions. Both pulses produce overlapping responses, indicating that under glutamate-only stimulation, receptor geometry has negligible influence on the paired-pulse kinetics. The singly glutamate-bound state [AGlu] (Fig. 5B) mirrors this behavior with two pronounced peaks—one after each pulse. The first peak arises within a fraction of a millisecond and reaches ∼ 15 receptors for all configurations. The second response, following the release at *t* = Δ, shows a comparable but slightly smaller amplitude. The doubly bound state [AGlu_2_] (Fig. 5C), exhibits a similar paired-pulse pattern but at larger amplitude. Each release event drives a near-synchronous rise of up to ∼ 45–50 receptors, followed by rapid decay as desensitization and unbinding proceed. Finally, the desensitized receptor population [*D*_0_] (Fig. 5D) integrates the outcome of both pulses. The two pulses trigger a gradual rise that peaks at ∼ 8–10 receptors, followed by a slow decay as receptors recover to the available pool.

**Figure 5:**
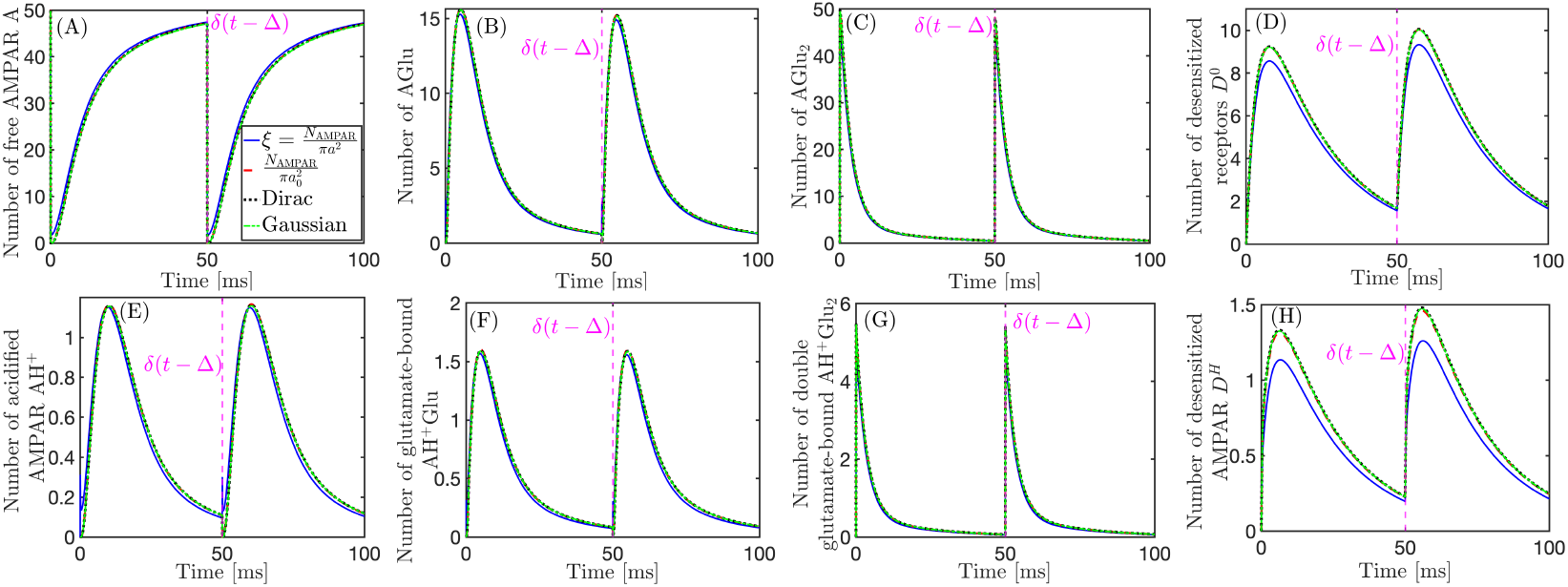
(A-D) Dynamics of AMPAR states under paired-pulse stimulation with glutamate release only (no protons). **(A)** Time course of free receptors [A]. **(B)** Singly glutamate-bound receptors [AGlu]. **(C)** Doubly glutamate-bound receptors [AGlu_2_]. **(D)** Desensitized receptors [*D*^0^]. Rate constants used in the simulation are *k*_−4_ = 200 s^−1^, *k*_−5_ = 200 s^−1^, *k*_des_ = 69.9 s^−1^, and *k*_rec_ = 64.1 s^−1^. 3000 glutamate were released to the cleft in the simulation. **(E-H) Dynamics of AMPAR states under paired-pulse stimulation with glutamate and proton release only (E)** Acidified receptors [AH^+^]. **(F)** Singly glutamate-bound acidified receptors [AH^+^Glu]. **(G)** Doubly glutamate-bound acidified receptors [AH^+^Glu_2_]. **(H)** Proton-driven desensitized receptors [*D*^*H*^]. Paired-pulse stimulation (Δ = 50 ms, is marked by the magenta dashed line *δ*(*t* − Δ)). Simulation parameters are listed in Table 2.1, with 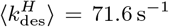 and 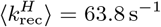 used for this figure. 3000 glutamate and 376 protons were released to the cleft in the simulation. Each curve compares four receptor spatial distributions: uniform full-disk (blue), uniform sub-disk (red dashed), clustered (black dotted), and broad Gaussian (green dash-dotted).

### 2.5 Paired–pulse response in the presence of protons

To assess how transient acidification influences receptor recovery between consecutive glutamate releases, we simulated paired-pulse activation at Δ = 50 ms while co-releasing protons (*N*_0,*H*_ = 376, baseline pH = 7.4) (Fig. 5E-H, Fig. S3A–C). Immediately after each pulse, the proton burst rapidly lowers the local pH from 7.4 to ∼ 4.5 near the release center, forming a short-lived acidification front that expands before dissipating within ∼ 0.5 ms (Fig. S3A). This transient wave locally elevates the desensitization rate and slows recovery (Fig. S3B–C), momentarily limiting receptor availability before the second event. Comparing the acidified and non-acidified branches, the free receptor population [AH^+^] (Fig. 5E) reaches only about 2–3% of the total [A] pool, indicating that most receptors remain non-protonated. Among glutamate-bound states, the single-bound acidified species [AH^+^Glu] constitutes ∼ 10% of its neutral counterpart [AGlu] (Fig. 5F), similarly to the double-bound form [AH^+^Glu_2_] (Fig. 5G). The proton-dependent desensitized population [*D*^*H*^] (Fig. 5H) consists ∼ 15% of [*D*^0^]. Direct comparison between the upper (glutamate-only) and lower (glutamate + proton) panels in Fig. 5 highlights the modulatory impact of transient acidification on AMPAR gating. Although the overall paired-pulse profiles remain qualitatively similar, the inclusion of protons systematically decreases the peak amplitudes of glutamate-bound states and desensitized receptors.

### 2.6 Dependence of paired–pulse responses on proton load

To examine how paired-pulse dynamics depend on proton load—and therefore on the local cleft pH—we analyzed the temporal evolution of receptor populations for increasing *N*_0,*H*_ values (10–1000) (Fig. 6A–F). As the proton load increases, acidification deepens (pH ≈ 6.0 → 4.0), progressively altering both activation and recovery phases.

**Figure 6:**
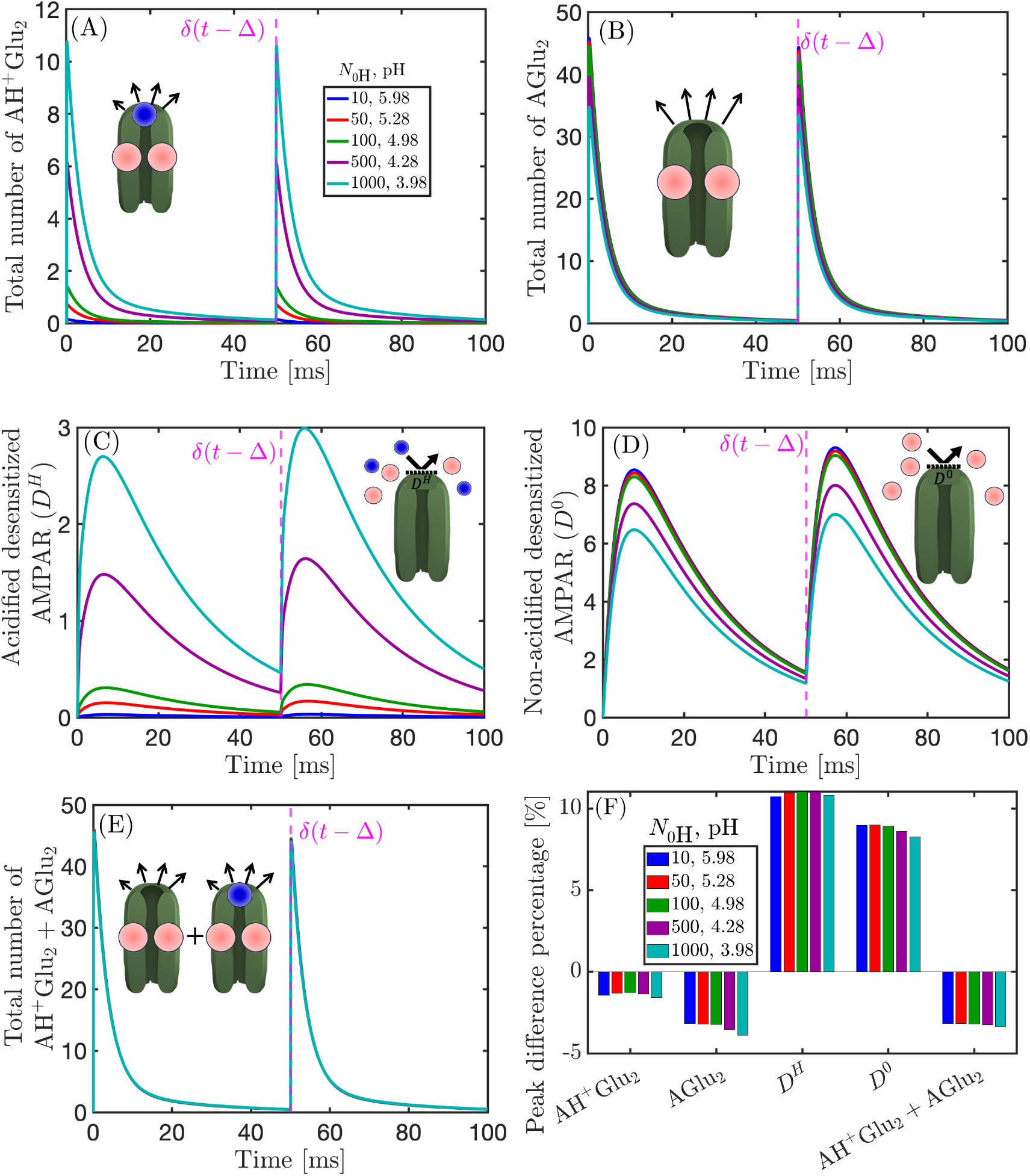
Responses of AMPAR species under paired-pulse stimulation with varying proton loads. **(A–E)** Temporal dynamics of the total number of receptors in each state for increasing proton release numbers *N*_0*H*_ = 10, 50, 100, 500, 1000, with corresponding cleft pH values shown in the legend. **(A)** Acidified double–glutamate bound receptors [AH^+^Glu_2_]. **(B)** Non–acidified double–glutamate bound receptors [AGlu_2_]. **(C)** Acidified desensitized state [D^*H*^]. **(D)** Non–acidified desensitized state [D^0^]. **(E)** Combined double–glutamate occupancy ([AH^+^Glu_2_] + [AGlu_2_]). **(F)** Percentage change in peak amplitude between the second and first responses for each species. The magenta dashed line indicates the inter-pulse interval Δ, marking the second stimulation as *δ*(*t*−Δ).

The acidified doubly glutamate-bound population [AH^+^Glu_2_] exhibits two rapid transients following the paired stimuli (Fig. 6(A)). For both the first and second pulse, the peak rises from ∼ 0 receptors at *N*_0,*H*_ = 10 to nearly ∼ 11 at *N*_0,*H*_ = 1000. The non-acidified counterpart [AGlu_2_] dominates the total activation (Fig. 6(B)), with peaks of ∼ 35 − 45 receptors, from high to low pH.

The acidified desensitized population [D^*H*^] accumulates gradually after the first release (Fig. 6(C)), increasing from ∼ 0 at *N*_0,*H*_ = 10 to ∼ 2.5 receptors at *N*_0,*H*_ = 1000. The non-acidified desensitized population [D^0^] (Fig. 6D) shows a similar two-pulse pattern but with higher overall occupancy, peaking around ∼ 6–8 receptors and exhibiting a mild increment between pulses.

The open-channel ensemble [AH^+^Glu_2_] + [AGlu_2_] exhibits comparable behavior across all conditions, with maxima around 45 receptors with only a slight decrease in the second pulse (Fig. 6(E)).

Quantitative comparison of the peak amplitudes between the two pulses (Fig. 6F) reveals that the acidified and non-acidified open states [AH^+^Glu_2_] and [AGlu_2_] undergo mild paired-pulse depression of ∼ 1–4 % across all proton loads. In contrast, the desensitized populations [*D*^*H*^] and [*D*^0^] display a facilitation of ∼ 6–10 %. The combined open-channel occupancy [AH^+^Glu_2_]+[AGlu_2_] shows a small negative shift of about ∼3 %, consistent across pH levels.

### 2.7 Dependence of paired–pulse responses on receptor recovery and desensitization rates

Following the results presented in Fig. 6F, where the summed open-state population [AH^+^Glu_2_]+[AGlu_2_] shows an apparently flat dependence of the peak-difference percentage on pH, we next sought to determine whether this apparent invariance persists across a wider kinetic range. Specifically, we analyzed the influence of the desensitization and recovery rates (*k*_des_ and *k*_rec_) on the paired–pulse response to test whether a kinetic threshold exists beyond which the flatness of the peak change across pH breaks. To this end, we systematically scanned the backward rates *k*_−*i*_ over the range 25:25:500 s^−1^, applying the constraint *k*_−1_ = *k*_−2_ = *k*_−3_ = *k*_−4_ = *k*_−5_ to minimize computational cost while retaining representative kinetic behavior. This reduction was necessary since a full parameter sweep across (*k*_−1_, *k*_−2_, *k*_−3_, *k*_−4_, *k*_−5_) ∈ *{*25, 50, 75, …, 500*}*^5^ would result in more than 3.2 *×* 10^6^ combinations.

The resulting two–dimensional modulation map (Fig. S4E) reveal that the surface of the second–pulse amplitude difference remains remarkably uniform across the physiological pH range (4–6), even when *k*_−*i*_ varies over nearly two orders of magnitude. For *k*_−*i*_ → 300 s^−1^, the percentage change saturates, indicating that receptor recovery becomes effectively instantaneous relative to the interpulse interval. The protonated open state [AH^+^Glu_2_] (Fig. S4A) exhibits a mild paired–pulse depression at slow kinetic regimes (*k*_−*i*_ *<* 100 s^−1^), with peak differences around 15 %. The non–protonated counterpart [AGlu_2_] (Fig. S4B) shows a similar trend but with larger absolute modulation, reaching up to ∼30 % decrease for the slowest rates, reflecting its higher sensitivity to recovery kinetics in the absence of proton stabilization. The desensitized populations display the complementary pattern. The protonated desensitized state [D^*H*^] (Fig. S4C) shows a clear increment of desensitized receptors between peaks, rising by up to ∼30 % when backward constant rates tend to ∼ 25 s^−1^ (lowest examined value)), whereas the neutral desensitized population [D^0^] (Fig. S4(D)) exhibits a milder rise of up to ∼14 %.

## 3 Discussion and conclusions

### 3.1 Synaptic acidification impact on AMPAR occupancy and recovery

To evaluate the impact of acidification on AMPAR, we develop here a coupled diffusion–reaction model: the model allows to explore the role of the release location, the size of the sub-disk from which glutamate molecules are released (Fig. 1D-F), and more important the unknown backward rate constants (Fig. 1H), showing that in a large range of these parameters, receptor acidification affects mildly (10-15% of bound receptors are acidified) the post-synaptic current. Acidification takes a portion of 10-15% from receptor desensitization and recovery dynamics, leading to longer recovery, which could be significant during multiple stimulations.

Proton clearance within the synaptic cleft is primarily dominated by diffusion occurring in less than 1 ms, independent of vesicular position. Such geometry-insensitive decay is consistent with earlier analyses of glutamate diffusion in synaptic microdomains, which showed that transmitter clearance is dominated by rapid escape and uptake at the cleft boundaries and glial organization [10, 27, 28].

Both proton and glutamate ligands act on sub-millisecond timescales (Fig. 2), however, the lifetime of protonated complexes is longer than that of glutamate-bound species by roughly a factor of 2 (1 ms vs 0.5 ms). The doubly bound acidified state [AH^+^Glu_2_] remains rare and reaches no more than 2% of the overall pool (Fig. 2D), as its formation requires a narrow temporal overlap between proton and glutamate binding before proton dissociation. In contrast, the non-protonated glutamate-bound states [AGlu] and [AGlu_2_] dominate the steady-state occupancy with up to 70% of the total pool (Fig. 2E-F). Single glutamate-only occupancy distribution is consistent with previous modeling [11], who reported that for a single vesicular release AMPARs are predominantly found in low-occupancy glutamate-bound states, with the two-glutamate configuration being the most probable and higher-order bindings (three- or four-glutamate) occurring only infrequently. Although native AMPARs are known to be tetrameric and can in principle bind up to four glutamate molecules, our kinetic scheme restricts binding to two glutamates per receptor.

### 3.2 pH-dependent reshaping of AMPAR state distributions

Varying the proton release number *N*_0_ reveals that stronger acidification selectively amplifies proton-dependent pathways while suppressing glutamate-driven ones (Fig. 3). As *N*_0_ increases from 10^2^ to 10^4^ (pH ≈ 5 to pH ≈ 3), maximum number of acidified populations such as [AH^+^] and [AH^+^Glu] increase by nearly an order of magnitude (from ∼ 5 to ∼ 30), whereas non-protonated states [AGlu] and [AGlu_2_] decrease correspondingly (from ∼ 30 to ∼ 5 and from ∼ 2 to ∼ 0.5, respectively). This reverse relationship arises as increasing the proton load drives a larger fraction of receptors into protonated configurations which compete with and replace the non-protonated glutamate-bound states, thereby reducing the pool of receptors available for purely glutamate-driven activation.

Since protonated AMPARs have a higher propensity to transition into non-conducting or desensitized modes, this shift in state occupancy naturally leads to a lower open probability and slower recovery from inactivation under acidic conditions, consistent with the observations [37, 38].

### 3.3 Proton-induced stabilization of desensitized states during paired-pulse stimulation

Under paired-pulse stimulation, transient acidification exerts a modest but consistent modulatory effect on AMPAR kinetics. When both glutamate and glutamate+proton is/are released, all receptor distributions yield nearly identical, overlapping depletion–recovery cycles, confirming that geometry plays a negligible role in short-term AMPAR PPR dynamics. When protons co-release, moderate changes emerge between the different AMPAR species and activation and desensitization. For the protonated scenario, the open-channel state ([AH^+^Glu_2_], consumes around 10% of the open state [AGlu_2_]), when protons are not introduced. When protons are introduced, around 15% of the desensitized population ([*D*^0^]), are transformed into [*D*^*H*^], suggesting an apparently stable overall recovery (Fig. 5). The paired-pulse traces reflect proton-induced stabilization of non-conducting receptor states and slower recovery from desensitization, consistent with [37, 38].

### 3.4 Limitations and future directions: incorporating AMPAR trafficking

Although the present formulation assumes static receptor positions, synapses exhibit continuous AMPAR trafficking within and around the postsynaptic density [54, 55]. Such lateral mobility and dynamic redistribution may influence how rapidly receptors replenish the pool available for activation during repetitive stimulation. While protonation profoundly modulates AMPAR gating [37, 38], its impact on receptor diffusion or nanoscale mobility has not yet been experimentally quantified, leaving the extent to which local acidification shapes trafficking-mediated recovery an open question.

Future extensions should incorporate receptor trafficking into the coupled reaction–diffusion–kinetic framework to reveal how nanoscale AMPAR exchange shapes synaptic reliability and short-term plasticity. One possible approach is to introduce simplified trafficking dynamics directly into the present simulation by allowing a defined fraction of receptors to be removed from the synapse after each stimulation event—mimicking endocytic loss or lateral escape from the PSD—while simultaneously introducing a corresponding population of newly available receptors. Iteratively applying this update between successive pulses would enable us to track how activity-dependent replenishment and depletion reshape the evolving receptor-state distribution, and thereby quantify the extent to which trafficking contributes to paired-pulse responses and recovery kinetics.

## 4 Method

The method is decomposed into several subsections: we first present a diffusion model of protonation. We then describe glutamate diffusion and interaction with AMPAR. Finally, we describe a model with glutamate and protons inside the synaptic cleft.

### 4.1 Reaction-diffusion modeling of AMPAR acidification during a single vesicular release event

We describe here the modeling of synaptic cleft acidification based on the proton dynamics. We approximate the motion of protons by a diffusion process. In addition, protons can interact with AMPARs, and eventually leave the synaptic cleft. The interaction between H^+^ and AMPARs is governed by the chemical kinetics,

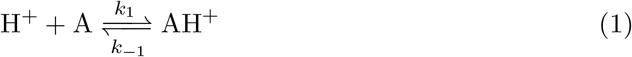

where AH^+^ is the bound state of AMPARs with a proton, and *k*_1_ and *k*_−1_ represent the association and dissociation rate constants, respectively. Thus, the molecular interaction at each receptor depends on the local proton concentration.

### 4.2 Synaptic Cleft Geometry Approximated by a flat cylinder

The synaptic cleft is approximated as a cylinder with a height *L* and a radius *a* as can be seen in Fig. 1E. The upper base of the cylinder represents the presynaptic membrane, while the lower base corresponds to the postsynaptic membrane, where AMPARs are distributed. Since protons disappear at the cylindrical walls, we will impose an absorbing boundary for the population density. However, at top and bottom bases, molecules cannot escape and thus we impose a reflective boundary condition [56]. We use the radial distance *r*, height *z*, and time *t* as variables.

### 4.3 Proton diffusion in the synaptic cleft

The spatial-temporal concentration *u*(*r, z, t*) of protons is computed from the diffusion equation. The time evolution of the proton concentration inside the cleft domain Ω, where the upper and lower synaptic surface ∂Ω_*r*_ [57] are reflective and protons can disappear at the cleft boundary ∂Ω_*s*_, is the solution of

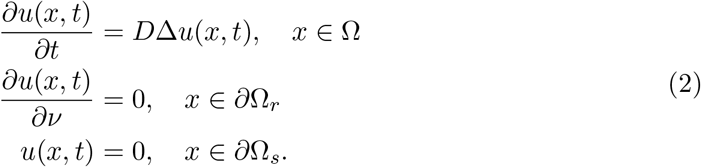

Proton release is modeled as an instantaneous point source at the top center of the cylinder (*r* = 0, *z* = *L*) (with a Dirac delta-function):

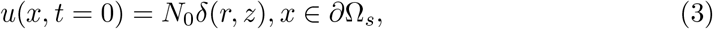

where *N*_0_ is the initial released protons. The solution (obtained by separation of variables [58, 59]) is

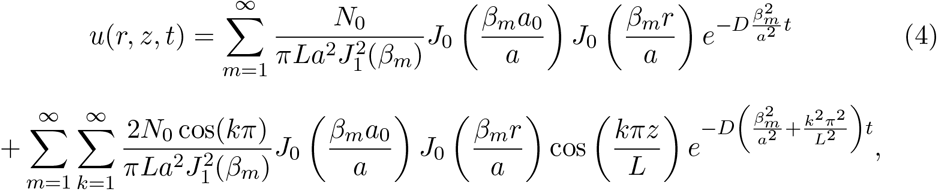

where *J*_0_(*x*) *J*_1_(*x*) are the zeroth and first order Bessel functions of the first kind, respectively, 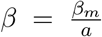, *m* = 1, 2, 3, …, where *β*_*m*_ are the roots of *J*_0_, and *a*_0_ is a sub-disk radius located on the pre-synaptic terminal from which the protons are uniformly released. At the surface of the post-synaptic cleft *z* = 0, the concentration is

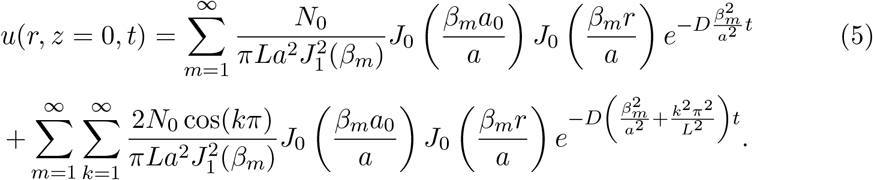

Finally, the surface concentration within the cleft is the average over the height given by

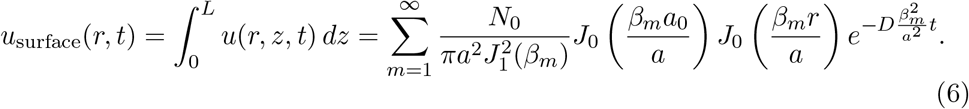

Finally, the total number of protons remaining in the cleft at time *t* is given by

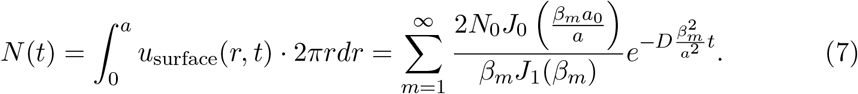

### 4.4 Modeling transient acidification of AMPAR

Acidified AMPA receptors are modeled by the chemical reaction in eq.(1), where the rate of arrival of H^+^ to a single AMPAR is diffusion limited. Using a radial distribution of AMPAR, eq.(1) can be simplified to

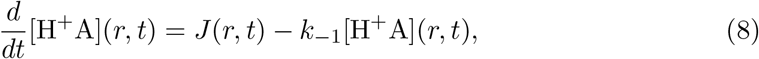

where the molecular flux is approximated as [58, 60]

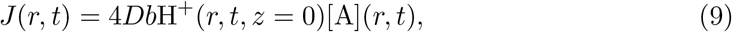

where [A](*r, t*) is the local receptor concentration in the annulus (*r, r* + *dr*) and *b* is the radius of an AMPA receptor. We shall consider four possible densities of AMPA receptors:1-uniform on the cleft, 2-uniform on a sub-disk, 3-located at the center of the post-synaptic cleft (Dirac distribution) and 4-Gaussian-like distribution, that is for *r* ≤ *a*

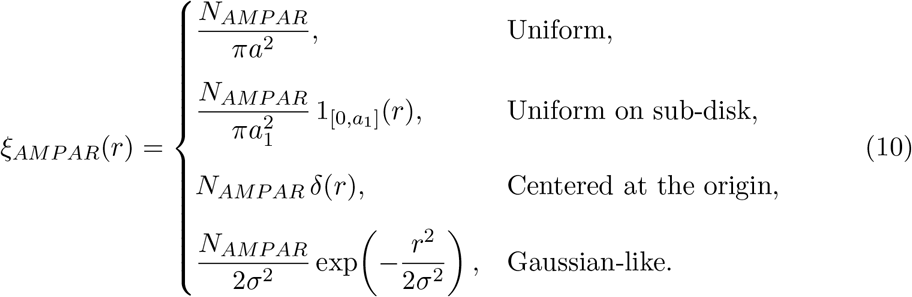

where 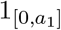 denotes the indicator function and total number of receptor is given by

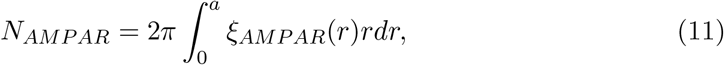

#### 4.4.1 Acidification kinetics equations

We derive here kinetics equations for acidification. Using the mass conservation of AMPAR, we get

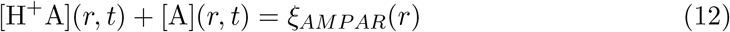

Thus the bound fraction *y*(*r, t*) = [H^+^A](*r, t*) of acidified receptor follows

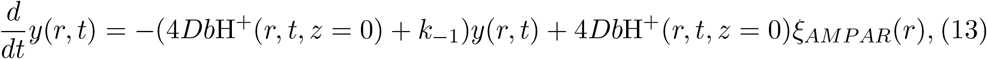

where *u*(*r, t*, 0) = H^+^(*r, t, z* = 0). A direct integration leads to

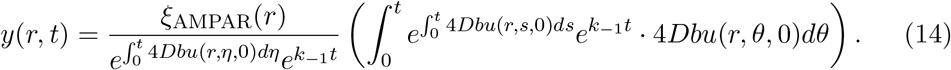

Thus, the dynamics of the total number of acidified AMPAR is given by

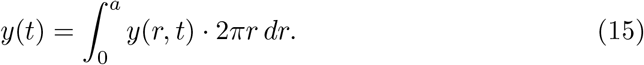

#### 4.4.2 Total number of acidified AMPAR following a single vesicular release

The total fraction of acidified AMPA receptors following a vesicular release is computed from eq.(1) by considering *k*_−1_ = 0 in the limit of *t* goes to infinity, leading to

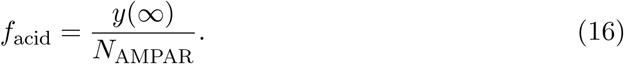

Thus

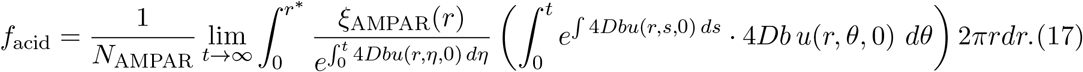

### 4.5 Modeling AMPAR activation with protons and glutamates

We now add glutamate in our modeling, on top of the proton dynamics within the synaptic cleft. Thus, AMPARs can bind glutamate, either in their protonated form (AH^+^) or without prior proton binding. The six states are given by A, AH^+^, AH^+^Glu, AH^+^Glu_2_, AGlu, AGlu_2_. The associated chemical reactions are:

1. Protonation of A to form AH^+^:

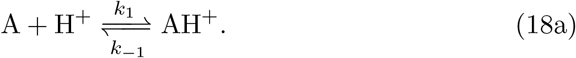
2. Complex formation between AH^+^ and the first glutamate molecule Glu:

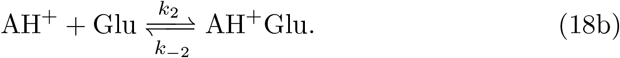
3. Complex formation between AH^+^Glu and the second glutamate molecule Glu:

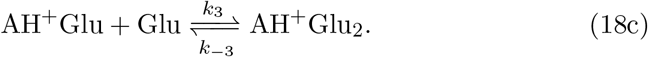
4. First glutamation of A to form AGlu:

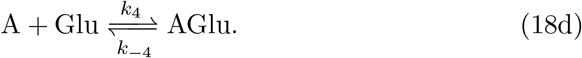
5. Second glutamation of AGlu to form AGlu_2_:

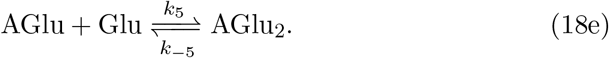

The parameters *k*_*i*_ and *k*_−*i*_ denote the forward and backward reaction rates, respectively.

#### 4.5.1 Glutamate Diffusion in the Cleft

Since glutamate is diffusing in the cleft, the concentration [Glu] = *u*_*G*_ satisfies eq.(2), where glutamate molecules are released instantaneously at a point source, at the top center of the cylinder (*r* = 0, *z* = *L*), as protons. Thus

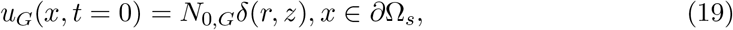

where *N*_0,*G*_ is the total initial of released glutamate molecules.

#### 4.5.2 Transition between species of AMPAR

Based on the kinetics eqs.(18a-18e), and eq.(9), we obtain the chemical differential equations for the protonated and glutamate-bound AMPARs

1. Protonation of A to form AH^+^:

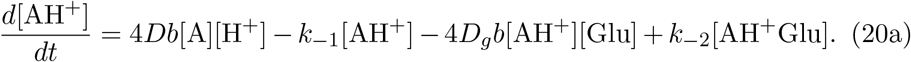
2. Complex formation between AH^+^ and Glu:

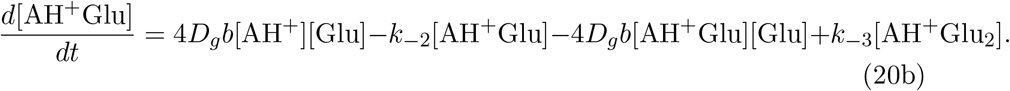
3. Complex formation between AH^+^Glu and Glu:

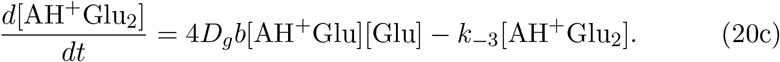
4. First glutamate binding of A to form AGlu:

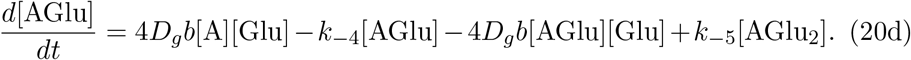
5. Second glutamate binding of AGlu to form AGlu_2_:

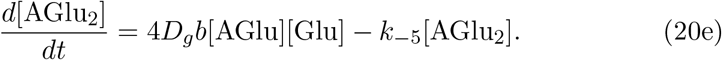

In this formulation, all forward binding rates *k*_*i*_ have been replaced by the corresponding diffusion-limited fluxes, 4*Db* for protons and 4*D*_*g*_*b* for glutamate, where *D*_*g*_ is the glutamate diffusion coefficient. Finally, the mass-conservation of AMPAR is given by

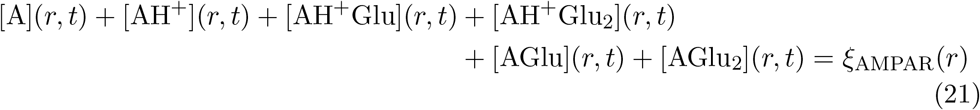

#### 4.5.3 Numerical implementation

We have simulated the concentrations *u*(*r, t*) and *u*_*g*_(*r, t*), solutions of the diffusion (eq.(2)) in cylindrical geometry, evaluated at the postsynaptic membrane plane (*z* = 0). These concentration fields were precomputed over a uniform time vector (*t* ∈ [4 *×* 10^−5^, 2 *×* 10^−3^]s, *n*_*t*_ = 1000, where *n*_*t*_ is the total number of time nodes) and radial grids matched to the receptor profiles.

Finally, we solve the kinetics described in the set of equations in (20) for the acidified only (AH), acidified and singly glutamate-bound (AHG), acidified and doubly glutamate-bound (AHG_2_), single glutamate-bound non-protonated (AGlu), and doubly glutamate-bound non-protonated (AGlu_2_). The unbound receptor population was determined by conservation of the local receptor density *ξ*_AMPAR_(*r*). The ODEs were integrated at each radial grid point using MATLAB’s solver *ode15s*, where the diffusion solutions *u*(*r, t*) and *u*_*g*_(*r, t*) served as time-dependent inputs. The resulting radial–temporal solutions for each species were then integrated over the postsynaptic disk using cylindrical quadrature to obtain the total number of receptors in each state as was shown in eq.(11).

#### 4.5.4 Receptor Distribution

To examine the role of receptor topography, we considered the four spatial receptor distributions described in eq.(10). Parameters are specified in table 2.1

### 4.6 Modeling protonation effect on AMPAR desensitization and recovery rate constants

Since AMPAR desensitization and recovery rate constants can vary with pH, we now add these two constants into our AMPAR-acidification-glutamate binding modeling. We thus present two different states: desensitized *D*^0^ (non-protonated) and *D*^*H*^ (protonated). The new rate equations become the following: For the non-protonated state, we use the following baseline rates

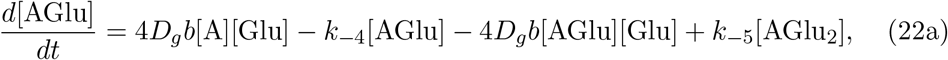

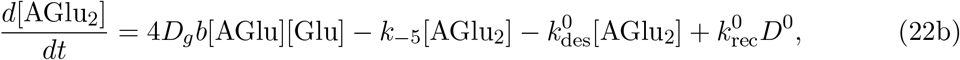

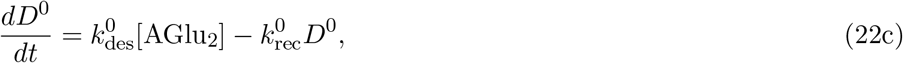

where 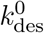 and 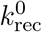 are the rate constants for the non-protonated state e.g. pH = 7.4. For the protonated state

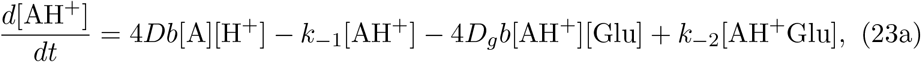

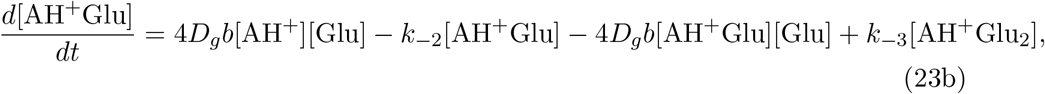

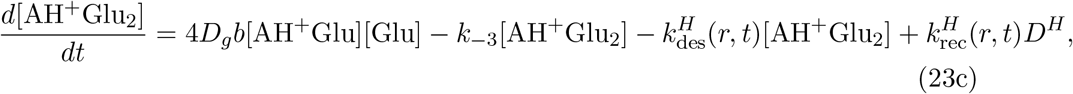

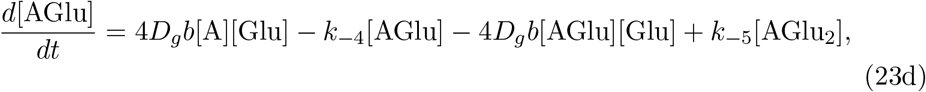

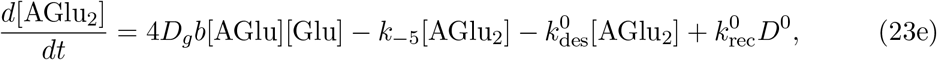

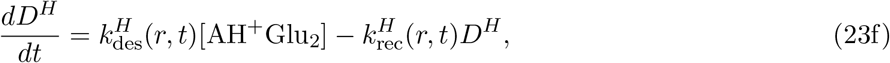

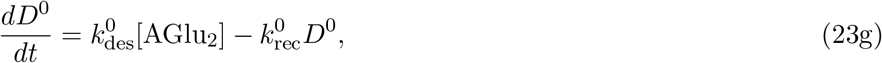

where we now linearly interpolate between the two pH values 5.5 and 7.4, leading

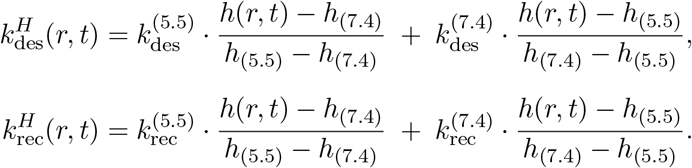

where

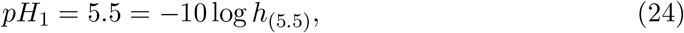

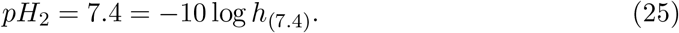

For the wild type of AMPAR which we model here, GluA2+TARP*γ*2, [37, 38] report (pH 7.4 vs pH 5.5): case A — desensitization time for a receptor which opens withoutprotons (AGlu_2_; pH 7.4 baseline): 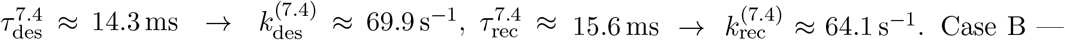 desensitization time for a receptor which opens without/with protons (AGlu_2_, AH^+^Glu_2_; acidic, pH 5.5): 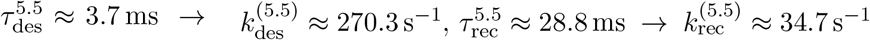.

Here we define how the proton–modulated rates are computed as a function of pH.

1. For the experimental window (5.5 ≤ *pH* ≤ 7.4): The rates are obtained by linear interpolation between the reference values at *pH* = 7.4 and *pH* = 5.5:

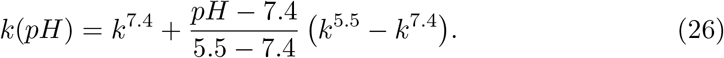
2. For strongly acidic conditions (*pH <* 5.5): The rates are extrapolated by extending the same straight line defined in the interval [5.5, 7.4]:

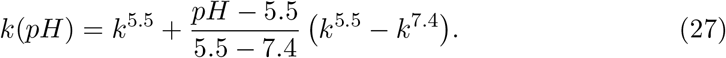

Applying this rule to both proton–dependent transitions:

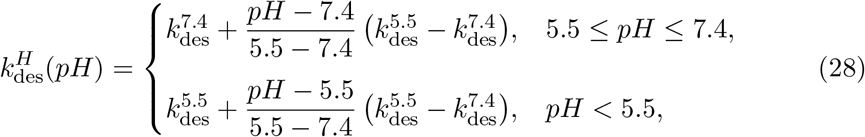

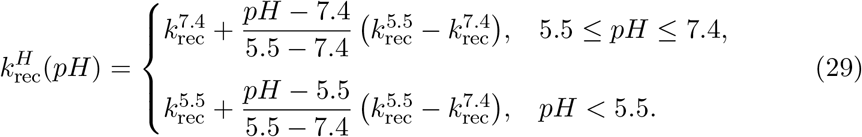

### 4.7 Modeling Paired-Pulse Ratio (PPR)

We have implemented a paired-pulse protocol consisting in releasing neurotransmitters separated by a time interval Δ = 50ms. We release both (H^+^) and glutamate, at *t* = 0 and *t* = Δ, with amplitudes 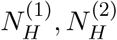 and 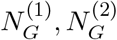, respectively, so that the concentrations are given by

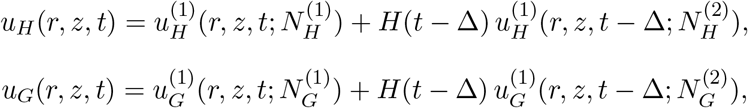

where *H*(·) is the Heaviside step function and *u*^(1)^ is the single-pulse solution Thus, the second pulse is incorporated simply as a time-shifted copy of the first.

## Conflict of interest

The authors declare no conflict of interest.

## Acknowledgments

D.H. research is supported by an ANR-23-CE46-0016 AnalysisSpectralEEG and AstroXcite and the European Research Council (ERC) under the European Union’s Horizon 2020 research and innovation program (grant agreement No 882673) and PoC 101212658.

## Supplementary Information

### Stating the Mathematical Problem for diffusion

The governing dimensional PDE is:

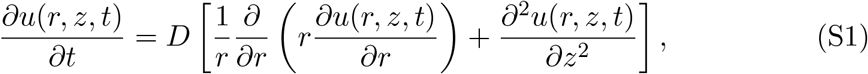

where *u*(*r, z, t*) is the concentration of particles, *D* is the diffusion coefficient, *r* is the radial coordinate and *z* is the axial coordinate. The domain is a cylinder of radius *a* and height *L*:

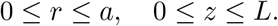

Boundary conditions:

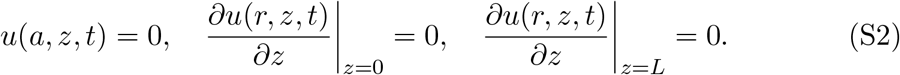

By using separation of variables method and assume a solution of the form *u*(*r, z, t*) = *R*(*r*)*Z*(*z*)*T* (*t*), we get the general solution

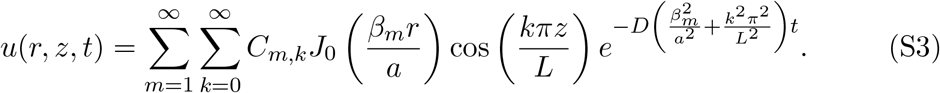

The initial condition is:

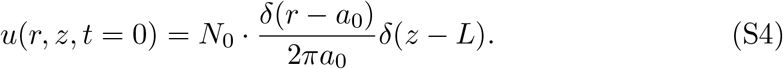

The coefficients *A*_*m,k*_ are obtained by projecting the initial condition onto the eigenfunctions, and we get the particular solution

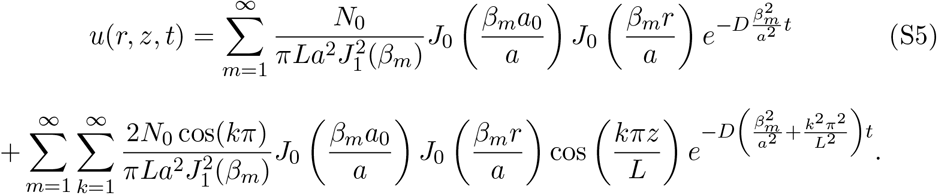

At the surface of the post synaptic cleft (*z* = 0), the dimensional concentration is:

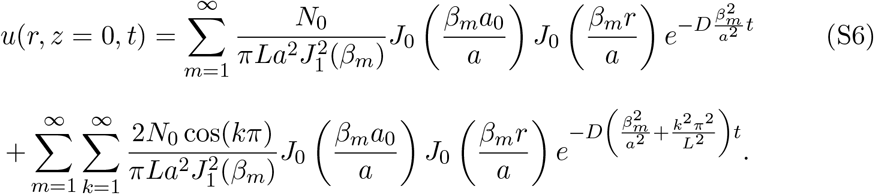

The total surface concentration is therefore

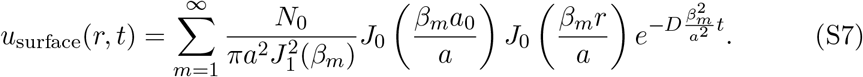

The spatio-temporal amount of protons is therefore

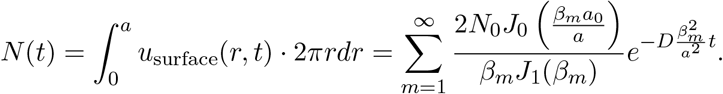

### Backward rate constants impact on the maximum of AMPAR-bound species

Given that the backward reaction rate constants *k*_−1_, *k*_−2_, *k*_−3_ (proton–related) and *k*_−4_, *k*_−5_ (glutamate–related) remain experimentally unconstrained, we performed a parameter–stability analysis to assess the robustness of the model across a broad kinetic range. This examination aimed to verify that the predicted receptor dynamics remain qualitatively consistent under variations of both proton and glutamate unbinding rates. We therefore scanned combinations of *k*_−*i*_ and evaluated the temporal evolution of all receptor states to identify regimes where the reaction system remains stable and physiologically meaningful (Fig. S1).

**Figure S1:**
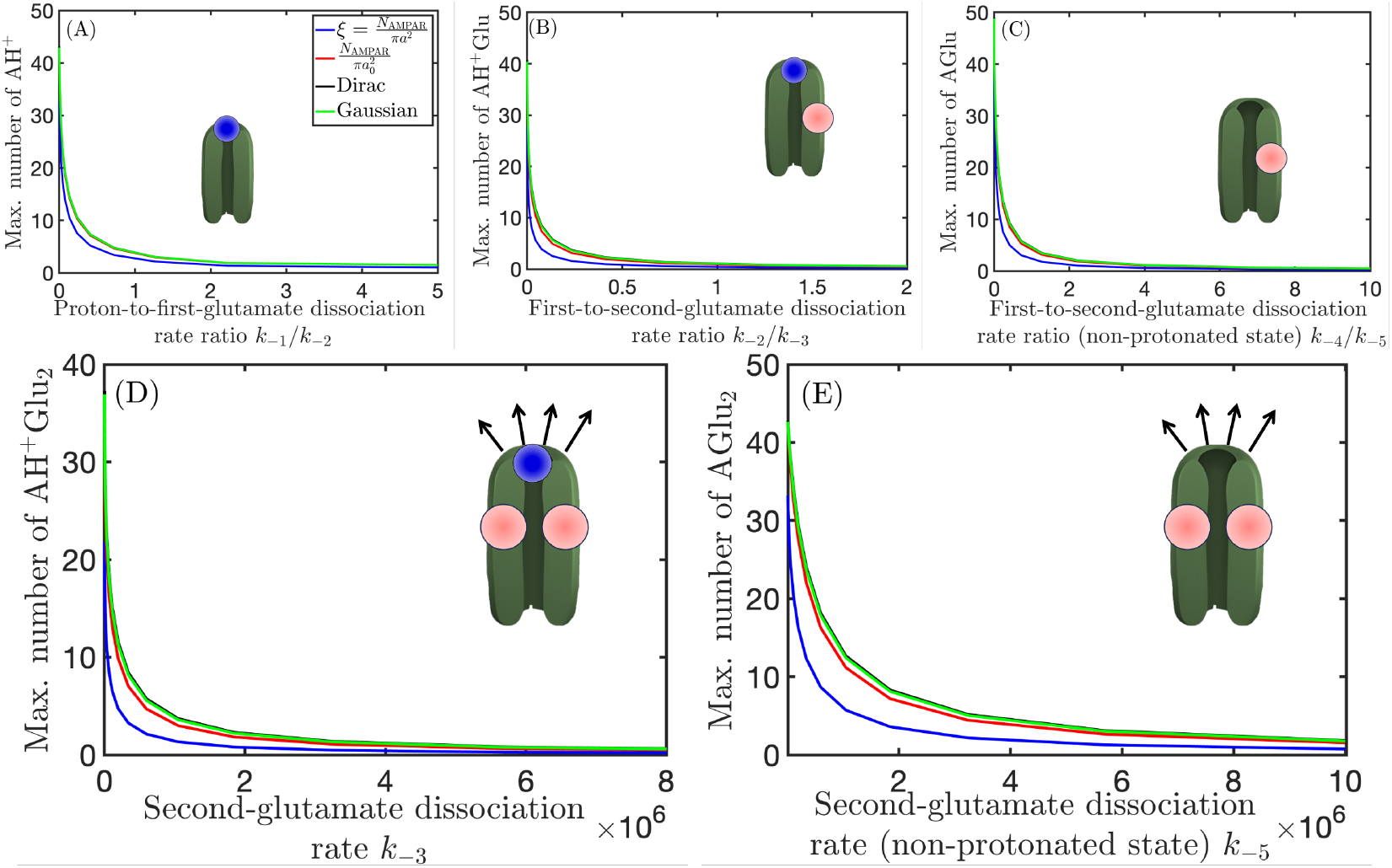
Maximum number of bound AMPARs as a function of reverse reaction rate ratios. **(A)** Protonated AMPARs [AH^+^] vs the ratio *k*_−1_*/k*_−2_. **(B)** Singly glutamate-bound acidified AMPARs [AH^+^Glu] vs *k*_−2_*/k*_−3_. **(C)** Singly glutamate-bound non-acidified AMPARs [AGlu] vs *k*_−4_*/k*_−5_. **(D)** Doubly glutamate-bound acidified AMPARs [AH^+^Glu_2_] vs *k*_−3_. **(E)** Doubly glutamate-bound non-acidified AMPARs [AGlu_2_] vs *k*_−5_. Each curve corresponds to one of the four receptor spatial distributions: uniform full-disk, uniform sub-disk, Dirac-like, and Gaussian.

### AMPAR-bound species relativity

To further characterize receptor dynamics in the absence of unbinding, we examined the limiting case where all backward reaction rate constants were set to zero (*k*_−1_ = *k*_−2_ = *k*_−3_ = *k*_−4_ = *k*_−5_ = 0). This test isolates the forward binding processes to evaluate the maximal extent of protonation and glutamate binding achievable under purely irreversible conditions. The analysis was performed across the four spatial receptor distributions *ξ*_AMPAR_(*r*), allowing comparison between uniform, sub-disk, Dirac-like, and Gaussian layouts (Fig. S2).

### Paired–pulse response in the presence of protons

To illustrate the immediate effect of proton release on the synaptic cleft environment, we simulated the spatiotemporal evolution of pH and its influence on the desensitization and recovery rate fields. This analysis quantifies how vesicular proton discharge produces a short-lived local acidification and transiently alters receptor kinetics prior to the second stimulus (Fig. S3).

**Figure S2:**
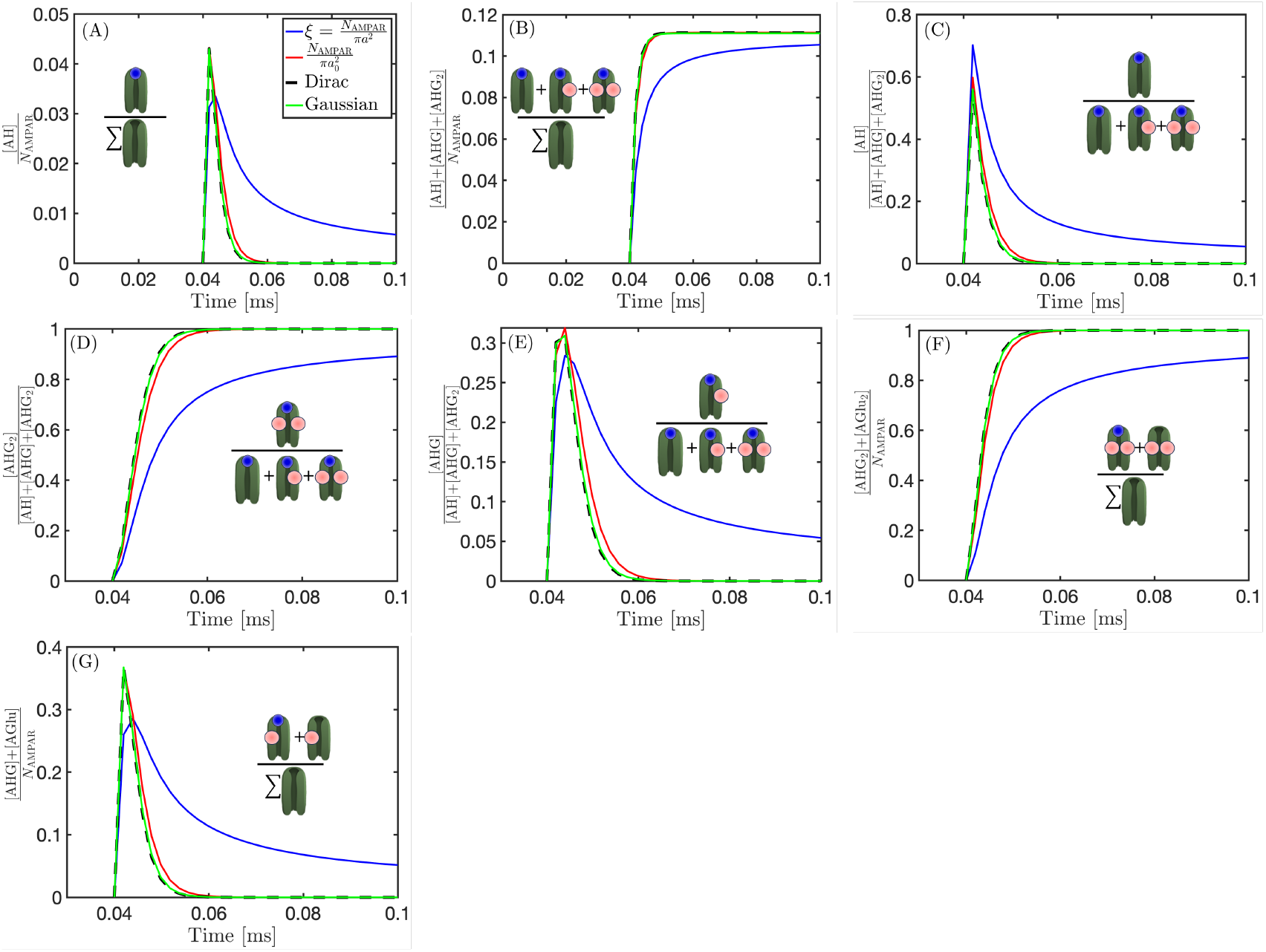
Normalized temporal fractions of AMPAR states for four spatial receptor distributions. All reverse (unbinding) reaction coefficients are set to zero. **(A)** Fraction of solely protonated receptors [AH^+^]*/N*_AMPAR_. **(B)** Fraction of total acidified receptors ([AH^+^] + [AH^+^Glu] + [AH^+^Glu_2_])*/N*_AMPAR_. **(C)** Fraction of the acidified pool that is protonated [AH^+^]*/*([AH^+^] + [AH^+^Glu] + [AH^+^Glu_2_]). **(D)** Fraction of the acidified pool in the open (doubly glutamate-bound) state [AH^+^Glu_2_]*/*([AH^+^] + [AH^+^Glu] + [AH^+^Glu_2_]). **(E)** Fraction of the acidified pool in the desensitized state [AH^+^Glu]*/*([AH^+^] + [AH^+^Glu] + [AH^+^Glu_2_]). **(F)** Fraction of all receptors in the open state ([AH^+^Glu_2_] + [AGlu_2_])*/N*_AMPAR_. **(G)** Fraction of all receptors in the desensitized state ([AH^+^Glu] + [AGlu])*/N*_AMPAR_. Each curve corresponds to one of the four receptor spatial distributions: uniform full-disk, uniform sub-disk, Dirac-like, and Gaussian.

**Figure S3:**
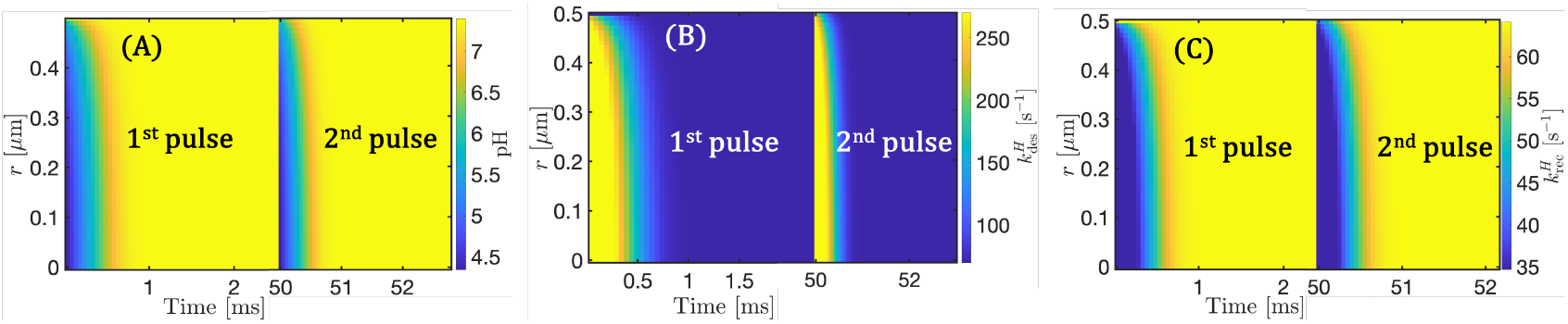
Paired-pulse response (PPR) in the presence of protons (baseline pH = 7.4, proton release *N*_0*H*_ = 376). **(A)** Spatiotemporal evolution of pH(*r, t*) across the post-synaptic cleft. **(B–C)** Spatial–temporal maps of the desensitization and recovery rates, 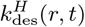 and 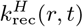, respectively.

### Dependence of paired–pulse responses on receptor recovery and desensitization rates

To explore the kinetic dependence of the paired–pulse response, we systematically varied the backward rate constants *k*_−*i*_ within the range 25−500 s^−1^ under equal–rate conditions (*k*_−1_ = *k*_−2_ = *k*_−3_ = *k*_−4_ = *k*_−5_). This analysis examines how receptor activation and desensitization balance across physiological pH values (4–6) as recovery rates accelerate (Fig. S4).

**Figure S4:**
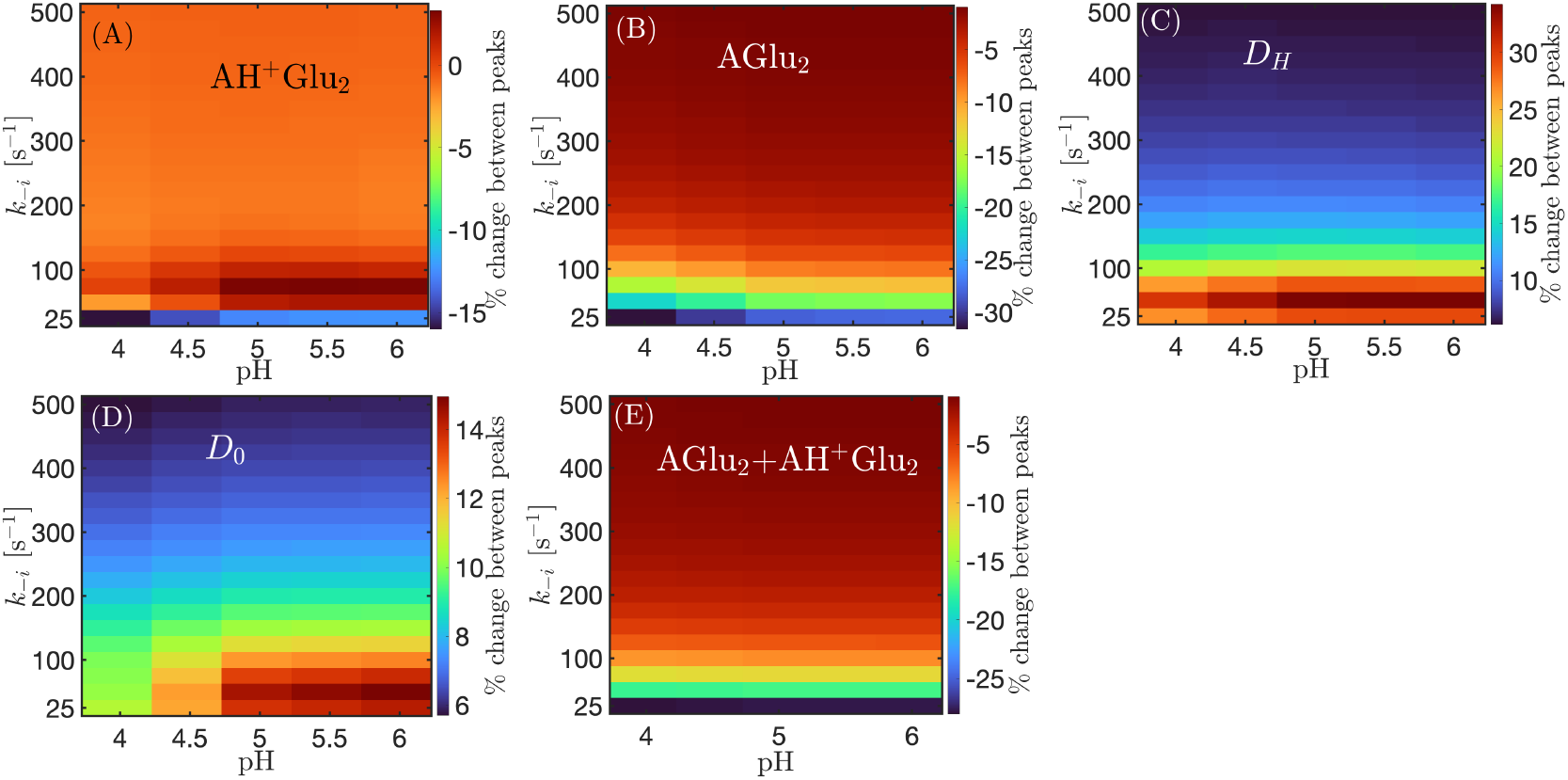
Dependence of paired–pulse responses on receptor recovery and desensitization rates. **(A–E)** Two–dimensional maps show the percentage change between the first and second peaks as a function of extracellular pH and recovery rate constant *k*_−*i*_ for five AMPAR–related species: **(A)** [AH^+^Glu_2_], **(B)** [AGlu_2_], **(C)** [*D*^*H*^], **(D)** [*D*^0^], and **(E)** total open–channel population [AH^+^Glu_2_] + [AGlu_2_].

## Notes

### Competing Interest Statement

The authors have declared no competing interest.

## References

[1] T. C. Südhof, “Neurotransmitter release: the last millisecond in the life of a synaptic vesicle,” Neuron, vol. 80, no. 3, pp. 675–690, 2013.

[2] B. Alberts, D. Bray, K. Hopkin, A. D. Johnson, J. Lewis, M. Raff, K. Roberts, and P. Walter, Essential cell biology. Garland Science, 2013.

[3] E. R. Kandel, J. H. Schwartz, T. M. Jessell, D. of Biochemistry, M. B. T. Jessell, S. Siegelbaum, and A. Hudspeth, Principles of neural science, vol. 4. McGraw-hill New York, 2000.

[4] K. B. Hansen, L. P. Wollmuth, D. Bowie, H. Furukawa, F. S. Menniti, A. I. Sobolevsky, G. T. Swanson, S. A. Swanger, I. H. Greger, T. Nakagawa, et al., “Structure, function, and pharmacology of glutamate receptor ion channels,” Pharmacological reviews, vol. 73, no. 4, pp. 1469–1658, 2021.

[5] R. A. Nicoll, “A brief history of long-term potentiation,” Neuron, vol. 93, no. 2, pp. 281–290, 2017.

[6] J. Lee, X. Chen, and R. A. Nicoll, “Synaptic memory survives molecular turnover,” Proceedings of the National Academy of Sciences, vol. 119, no. 42, p. e2211572119, 2022.

[7] S. Tomita, “Regulation of ionotropic glutamate receptors by their auxiliary subunits,” Physiology, vol. 25, no. 1, pp. 41–49, 2010.

[8] T. V. Bliss and G. L. Collingridge, “A synaptic model of memory: long-term potentiation in the hippocampus,” Nature, vol. 361, no. 6407, pp. 31–39, 1993.

[9] A. Nowacka, A. M. Getz, D. Bessa-Neto, and D. Choquet, “Activity-dependent diffusion trapping of ampa receptors as a key step for expression of early ltp,” Philosophical Transactions B, vol. 379, no. 1906, p. 20230220, 2024.

[10] D. Freche, U. Pannasch, N. Rouach, and D. Holcman, “Synapse geometry and receptor dynamics modulate synaptic strength,” PloS one, vol. 6, no. 10, p. e25122, 2011.

[11] A. Taflia and D. Holcman, “Estimating the synaptic current in a multiconductance ampa receptor model,” Biophysical journal, vol. 101, no. 4, pp. 781–792, 2011.

[12] D. A. Rusakov, L. P. Savtchenko, K. Zheng, and J. M. Henley, “Shaping the synaptic signal: molecular mobility inside and outside the cleft,” Trends in neurosciences, vol. 34, no. 7, pp. 359–369, 2011.

[13] N. Scheefhals and H. D. MacGillavry, “Functional organization of postsynaptic glutamate receptors,” Molecular and Cellular Neuroscience, vol. 91, pp. 82–94, 2018.

[14] T. E. Chater and Y. Goda, “The role of ampa receptors in postsynaptic mechanisms of synaptic plasticity,” Frontiers in cellular neuroscience, vol. 8, p. 401, 2014.

[15] C.A.M. de León-López, M. Carretero-Rey, and Z. U. Khan, “Ampa receptors in synaptic plasticity, memory function, and brain diseases,” Cellular and Molecular Neurobiology, vol. 45, no. 1, p. 14, 2025.

[16] K. M. Franks, T. M. Bartol, and T. J. Sejnowski, “A monte carlo model reveals independent signaling at central glutamatergic synapses,” Biophysical journal, vol. 83, no. 5, pp. 2333–2348, 2002.

[17] D. Choquet, “Linking nanoscale dynamics of ampa receptor organization to plasticity of excitatory synapses and learning,” Journal of Neuroscience, vol. 38, no. 44, pp. 9318–9329, 2018.

[18] T. Biederer, P. S. Kaeser, and T. A. Blanpied, “Transcellular nanoalignment of synaptic function,” Neuron, vol. 96, no. 3, pp. 680–696, 2017.

[19] M. Heine and D. Holcman, “Asymmetry between pre-and postsynaptic transient nanodomains shapes neuronal communication,” Trends in Neurosciences, 2020.

[20] D. A. Rusakov, “The role of perisynaptic glial sheaths in glutamate spillover and extracellular ca 2+ depletion,” Biophysical Journal, vol. 81, no. 4, pp. 1947– 1959, 2001.

[21] R. S. Zucker, “Minis: whence and wherefore?,” Neuron, vol. 45, no. 4, pp. 482– 484, 2005.

[22] G. M. Elias and R. A. Nicoll, “Synaptic trafficking of glutamate receptors by maguk scaffolding proteins,” Trends in cell biology, vol. 17, no. 7, pp. 343–352, 2007.

[23] T. A. Nielsen, D. A. DiGregorio, and R. A. Silver, “Modulation of glutamate mobility reveals the mechanism underlying slow-rising ampar epscs and the diffusion coefficient in the synaptic cleft,” Neuron, vol. 42, no. 5, pp. 757–771, 2004.

[24] C. Wichmann and T. Kuner, “Heterogeneity of glutamatergic synapses: cellular mechanisms and network consequences,” Physiological reviews, vol. 102, no. 1, pp. 269–318, 2022.

[25] X. Li et al., “Computational modeling of trans-synaptic nanocolumns and their influence on ampar currents,” Frontiers in Computational Neuroscience, 2022.

[26] E. Saftenku, “Modeling of slow glutamate diffusion and ampa receptor activation in the cerebellar glomerulus,” Journal of theoretical biology, vol. 234, no. 3, pp. 363–382, 2005.

[27] U. Pannasch, D. Freche, G. Dallérac, G. Ghézali, C. Escartin, P. Ezan, M. Cohen-Salmon, K. Benchenane, V. Abudara, A. Dufour, et al., “Connexin 30 sets synaptic strength by controlling astroglial synapse invasion,” Nature neuroscience, vol. 17, no. 4, pp. 549–558, 2014.

[28] L. P. Savtchenko and D. A. Rusakov, “Glutamate–transporter unbinding in probabilistic synaptic environment facilitates activation of distant nmda receptors,” Cells, vol. 12, no. 12, p. 1610, 2023.

[29] K. Zheng, A. Scimemi, and D. A. Rusakov, “Receptor actions of synaptically released glutamate: the role of transporters on the scale from nanometers to microns,” Biophysical journal, vol. 95, no. 10, pp. 4584–4596, 2008.

[30] F. Ventriglia and V. Di Maio, “Glutamate–ampar interaction in a model of synaptic transmission,” Brain research, vol. 1536, pp. 168–176, 2013.

[31] D. Choquet and A. Triller, “The role of receptor diffusion in the organization of the postsynaptic membrane,” Nature Reviews Neuroscience, vol. 4, no. 4, p. 251, 2003.

[32] K. Czöndör et al., “Unified quantitative model of ampa receptor trafficking at synapses,” Proc Natl Acad Sci USA, 2012.

[33] X. Xie, J.-S. Liaw, M. Baudry, and T. W. Berger, “Novel expression mechanism for synaptic potentiation: alignment of presynaptic release site and postsynaptic receptor,” Proceedings of the National Academy of Sciences, vol. 94, no. 13, pp. 6983–6988, 1997.

[34] A.-H. Tang, H. Chen, T. P. Li, S. R. Metzbower, H. D. MacGillavry, and T. A. Blanpied, “A trans-synaptic nanocolumn aligns neurotransmitter release to receptors,” Nature, vol. 536, no. 7615, pp. 210–214, 2016.

[35] M. Heine, L. Groc, R. Frischknecht, J.-C. Béïque, B. Lounis, G. Rumbaugh, R. L. Huganir, L. Cognet, and D. Choquet, “Surface mobility of postsynaptic ampars tunes synaptic transmission,” Science, vol. 320, no. 5873, pp. 201–205, 2008.

[36] I. Stockwell, J. F. Watson, and I. H. Greger, “Tuning synaptic strength by regulation of ampa glutamate receptor localization,” Bioessays, vol. 46, no. 7, p. 2400006, 2024.

[37] J. Ivica, N. Kejzar, H. Ho, I. Stockwell, V. Kuchtiak, A. M. Scrutton, T. Nakagawa, and I. H. Greger, “Proton-triggered rearrangement of the ampa receptor n-terminal domains impacts receptor kinetics and synaptic localization,” Nature Structural & Molecular Biology, vol. 31, no. 10, pp. 1601–1613, 2024.

[38] A. H. Larsen, A. M. Perozzo, P. C. Biggin, D. Bowie, and J. S. Kastrup, “Recovery from desensitization in glua2 ampa receptors is affected by a single mutation in the n-terminal domain interface,” Journal of Biological Chemistry, vol. 300, no. 3, p. 105717, 2024.

[39] R. H. Edwards, “The neurotransmitter cycle and quantal size,” Neuron, vol. 55, no. 6, pp. 835–858, 2007.

[40] G. Miesenböck, D. A. De Angelis, and J. E. Rothman, “Visualizing secretion and synaptic transmission with ph-sensitive green fluorescent proteins,” Nature, vol. 394, no. 6689, pp. 192–195, 1998.

[41] S. Cho and H. Von Gersdorff, “Proton-mediated block of ca2+ channels during multivesicular release regulates short-term plasticity at an auditory hair cell synapse,” Journal of Neuroscience, vol. 34, no. 48, pp. 15877–15887, 2014.

[42] M. Chesler, “Regulation and modulation of ph in the brain,” Physiological reviews, vol. 83, no. 4, pp. 1183–1221, 2003.

[43] S. H. DeVries, “Exocytosed protons feedback to suppress the ca2+ current in mammalian cone photoreceptors,” Neuron, vol. 32, no. 6, pp. 1107–1117, 2001.

[44] C.-K. Tong, K. Chen, and M. Chesler, “Kinetics of activity-evoked ph transients and extracellular ph buffering in rat hippocampal slices,” Journal of neurophysiology, vol. 95, no. 6, pp. 3686–3697, 2006.

[45] D. Zhang, J. Ivica, J. M. Krieger, H. Ho, K. Yamashita, I. Stockwell, R. Baradaran, O. Cais, and I. H. Greger, “Structural mobility tunes signalling of the glua1 ampa glutamate receptor,” Nature, vol. 621, no. 7980, pp. 877–882, 2023.

[46] E. C. Ihle and D. K. Patneau, “Modulation of α-amino-3-hydroxy-5-methyl-4-isoxazolepropionic acid receptor desensitization by extracellular protons,” Molecular pharmacology, vol. 58, no. 6, pp. 1204–1212, 2000.

[47] S. Lei, B. A. Orser, G. R. Thatcher, J. N. Reynolds, and J. F. MacDonald, “Positive allosteric modulators of ampa receptors reduce proton-induced receptor desensitization in rat hippocampal neurons,” Journal of Neurophysiology, vol. 85, no. 5, pp. 2030–2038, 2001.

[48] M. Koike, S. Tsukada, K. Tsuzuki, H. Kijima, and S. Ozawa, “Regulation of kinetic properties of glur2 ampa receptor channels by alternative splicing,” Journal of Neuroscience, vol. 20, no. 6, pp. 2166–2174, 2000.

[49] T. L. Stincic and M. E. Frerking, “Different ampa receptor subtypes mediate the distinct kinetic components of a biphasic epsc in hippocampal interneurons,” Frontiers in Synaptic Neuroscience, vol. 7, p. 7, 2015.

[50] A. L. Carbone and A. J. Plested, “Coupled control of desensitization and gating by the ligand binding domain of glutamate receptors,” Neuron, vol. 74, no. 5, pp. 845–857, 2012.

[51] S. R. Platt, “The role of glutamate in central nervous system health and disease– a review,” The Veterinary Journal, vol. 173, no. 2, pp. 278–286, 2007.

[52] L. P. Savtchenko and D. A. Rusakov, “The optimal height of the synaptic cleft,” Proceedings of the National Academy of Sciences, vol. 104, no. 6, pp. 1823–1828, 2007.

[53] D. Atasoy, M. Ertunc, K. L. Moulder, J. Blackwell, C. Chung, J. Su, and E. T. Kavalali, “Spontaneous and evoked glutamate release activates two populations of nmda receptors with limited overlap,” Journal of Neuroscience, vol. 28, no. 40, pp. 10151–10166, 2008.

[54] W.-K. Cho, N. Jayanth, B. P. English, T. Inoue, J. O. Andrews, W. Conway, J. B. Grimm, J.-H. Spille, L. D. Lavis, T. Lionnet, et al., “Rna polymerase ii cluster dynamics predict mrna output in living cells,” Elife, vol. 5, p. e13617, 2016.

[55] N. W. Cho, R. L. Dilley, M. A. Lampson, and R. A. Greenberg, “Interchromosomal homology searches drive directional ALT telomere movement and synapsis.,” Cell, vol. 159, pp. 108–21, Sept. 2014.

[56] Z. Schuss, Brownian dynamics at boundaries and interfaces. Springer, 2015.

[57] D. Holcman and Z. Schuss, Asymptotics of Elliptic and Parabolic PDEs. Springer, 2018.

[58] J. Crank, The mathematics of diffusion. Oxford university press, 1979.

[59] D. Holcman and Z. Schuss, “Stochastic narrow escape in molecular and cellular biology,” Analysis and Applications. Springer, New York, vol. 48, pp. 108–112, 2015.

[60] A. Singer, Z. Schuss, and D. Holcman, “Narrow escape and leakage of brownian particles,” Physical Review E, vol. 78, no. 5, 2008.

